# Pharmacologically targeting KRAS^G12D^ in PDAC models: tumor cell intrinsic and extrinsic impact

**DOI:** 10.1101/2023.03.18.533261

**Authors:** Vishnu Kumarasamy, Costakis Frangou, Jianxin Wang, Yin Wan, Andrew Dynka, Hanna Rosenheck, Prasenjit Dey, Ethan V. Abel, Erik S. Knudsen, Agnieszka K. Witkiewicz

## Abstract

Pancreatic ductal adenocarcinoma (PDAC) is an aggressive disease for which new therapeutic interventions are needed. Here we assessed the cellular response to pharmacological KRAS inhibition, which target the central oncogenic factor in PDAC. In a panel of PDAC cell lines, pharmaceutical inhibition of KRAS^G12D^ allele, with MRTX1133 yields variable efficacy in the suppression of cell growth and downstream gene expression programs in 2D culture. CRISPR screens identify new drivers for enhanced therapeutic response that regulate focal adhesion and signaling cascades, which were confirmed by gene specific knockdowns and combinatorial drug synergy. Interestingly, MRTX1133 is considerably more efficacious in the context of 3D cell cultures and *in vivo* PDAC patient-derived xenografts. In syngeneic models, KRAS^G12D^ inhibition elicits potent tumor regression that did not occur in immune-deficient hosts. Digital spatial profiling on tumor tissues indicates that MRTX1133 activates interferon-γ signaling and induces antigen presentation that modulate the tumor microenvironment. Further investigation on the immunological response using single cell sequencing and multispectral imaging reveals that tumor regression is associated with suppression of neutrophils and influx of effector CD8^+^ T-cells. Thus, both tumor cell intrinsic and extrinsic events contribute to response and credential KRAS^G12D^ inhibition as promising strategy for a large percentage of PDAC tumors.

## Introduction

In spite of significant effort, pancreatic ductal adenocarcinoma (PDAC) remains a largely-therapy recalcitrant disease for which overall five-year survival remains ∼11%^1–4^. Genetic analyses indicate that PDAC tumors harbor multiple high-potency oncogenic and tumor suppressive lesions^5–7^. While a myriad of targeted therapeutic approaches have been tested in clinical trials, in many contexts rationally targeted therapies exploiting genetic features of PDAC have failed to exhibit superiority to chemotherapy. For example, MEK inhibitors which would be expected to have broad efficacy in KRAS-driven PDAC have shown little efficacy^8^. Thus, developing new therapeutic strategies which target the genetics of PDAC remain a largely unrealized opportunity.

PDAC is dominated by oncogenic mutation of KRAS occurring in >95% of tumors^9^. Mutant KRAS acts through multiple signaling pathways that contribute to deregulated proliferation, invasion, and metastasis^10, 11^. The signaling mediated by mutant KRAS initiates at the membrane and is transduced by multiple effectors that represent independent drug targets. In PDAC cell lines, there is a variable response to KRAS depletion which generally mediates a cytostatic response^11, 12^. However, the genetic ablation of KRAS can be tolerated and such genetically modified PDAC cell lines can develop into tumors in xenograft models^13^. The targeted deletion of KRAS in GEMM models can lead to tumor regression; however, KRAS-independent tumors can evolve, with acquired resistance associated with the activation of bypass pathways (e.g. EGFR, AKT, YAP) that abrogate the dependence^14–16^.

The tumor microenvironment is particularly relevant in PDAC where the stromal compartment can contribute to the bulk of the tumor volume^17, 18^. This stroma has been proposed to both serve tumor inhibitory and promoting functions^19, 20^. In particular, it has been believed that the stroma contributes to exclusion of functional anti-tumor immune functions, and contributes to the lack of efficacy of immune checkpoint inhibitors and other forms of immunotherapy^4, 21^. Studies with the genetic modulation of KRAS have illustrated that tumor-mediated events contribute to broad ranging effects on the tumor microenvironment^22^.

KRAS inhibitors have been developed to the stage of FDA-approval in the context of agents selectively directed against the G12C mutant allele^23^. This mutation of KRAS is present at relatively low frequency (1-2%) in PDAC^10^. However, there is evidence that a G12C inhibitor (ie. Sotorasib) can have clinical activity in PDAC^24^, although there are clearly distinct mechanisms through which resistance can emerge^25, 26^ The most prevalent KRAS mutation in PDAC is G12D^10, 27^. Recently the agent MRTX1133 was described as a highly-selective and potent inhibitor of G12D, which has been shown to have efficacy in PDAC models^28, 29^. Here we explored the functional response to G12D inhibition in a spectrum of models that define features of sensitivity and resistance to this agent and underscore the importance of the tumor microenvironment on therapeutic efficacy.

## Results

### Heterogeneous cellular response to MRTX1133 in PDAC models

To define the impact of a pharmacological KRAS inhibitor, MRTX1133 we employed a panel of KRAS^G12D^ mutant PDAC cell lines that include both established and patient derived models ^45^. Following the treatment with MRTX1133 we observed a highly varied response based on the proliferation of the cells as monitored by live cell imaging. The established PDAC cell lines, ASPC1 and HPAF-II and a patient derived cell line, 828, were sensitive to MRTX1133 and displayed a robust cytostatic response (Fig. 1A). Interestingly, the other patient derived PDAC cell lines, 519, 1222 and 3226 were largely resistant, as cellular proliferation continued despite the presence of MRTX1133 (Fig.1A & S1). The EC50 values for MRTX1133 in the sensitive models were at the range of 25-50 nM, while in the resistant models, the EC50 is greater than 1 µM (Fig. 1B). To validate the specificity of MRTX1133 to KRAS^G12D^ mutants, we evaluated the anti-proliferative effect of this drug in KRAS^G12C^ mutant cell lines MiaPaCa2 and a patient derived PDAC cell line, UM53 (Fig. S1). As a control, we included the KRAS^G12C^ inhibitor, MRTX849. Based on the growth curves, the proliferation of MiaPaCa2 cells was robustly inhibited by MRTX849, whereas the MRTX1133 had no effect, confirming that the KRAS^G12D^ is its intracellular target (Fig. S1). Although, in UM53 cell line, the response to MRTX849 is modest, there is a significant difference in response as compared to MRTX1133 (Fig. S1). To interrogate how different cellular mechanisms are impacted by MRTX1133 to mediate an anti-proliferative effect, transcriptome analysis was carried out in different PDAC cell lines in the presence and absence of the KRAS inhibitor. Based on differential gene expression analysis the sensitive models, ASPC1, HPAF-II and 828 displayed a relatively greater number of genes that were differentially expressed in the presence of MRTX1133 as compared to the resistant models 519, 827 and 1229 indicating that the resistant models are essentially devoid of a transcription output downstream from KRAS inhibition (Fig.1C & D). Gene set enrichment analysis (GSEA) on the cell lines indicated that the downregulated genes are associated with E2F targets, MYC targets, G2/M check point and MTOR signaling (Fig. 1E). However, the magnitude of repression differs among the models, which correlates with their sensitivity to MRTX1133 (Fig. 1F). Since, the E2F target genes are known to regulate cell cycle progression via RB inactivation, biochemical analysis was performed in the PDAC cell lines following the treatment with MRTX1133. In the sensitive models, 828 and ASPC1 RB phosphorylation was significantly inhibited, which further resulted in the suppression of cyclin A, which is an E2F target (Fig. 1G). As expected, in the resistant models, 3226 and 519, RB remained phosphorylated with modest impact on cyclin A, indicating that cell cycle remains ongoing in the presence of the KRAS inhibitor (Fig. 1G). Consistent with the biochemical data, MRTX1133 significantly inhibited BrdU incorporation in the sensitive models confirming a robust cell cycle arrest, while moderate or no effect was observed in the resistant models (Fig. S1). Interestingly, the phosphorylation of ERK was inhibited across the PDAC models irrespective of their sensitivities to MRTX1133, indicating that at least this downstream pathway of KRAS is inhibited. However, this was not sufficient to inhibit RB phosphorylation and expression of downstream proteins (e.g. Cyclin A) in the resistant models (Fig. 1G). To confirm the inhibition of ERK phosphorylation by MRTX1133 is a consequence of KRAS^G12D^ inhibition, we compared its effect with MRTX849 on the MiaPaCa2 cells. Our data confirmed that inhibition of ERK phosphorylation is not affected by MRTX1133 (Fig. S1). In conclusion, although different PDAC models harbor oncogenic KRAS G12D mutation, there is heterogeneity in response to the pharmacological inhibitor, which differentially impacts on cell cycle.

**Figure 1.**
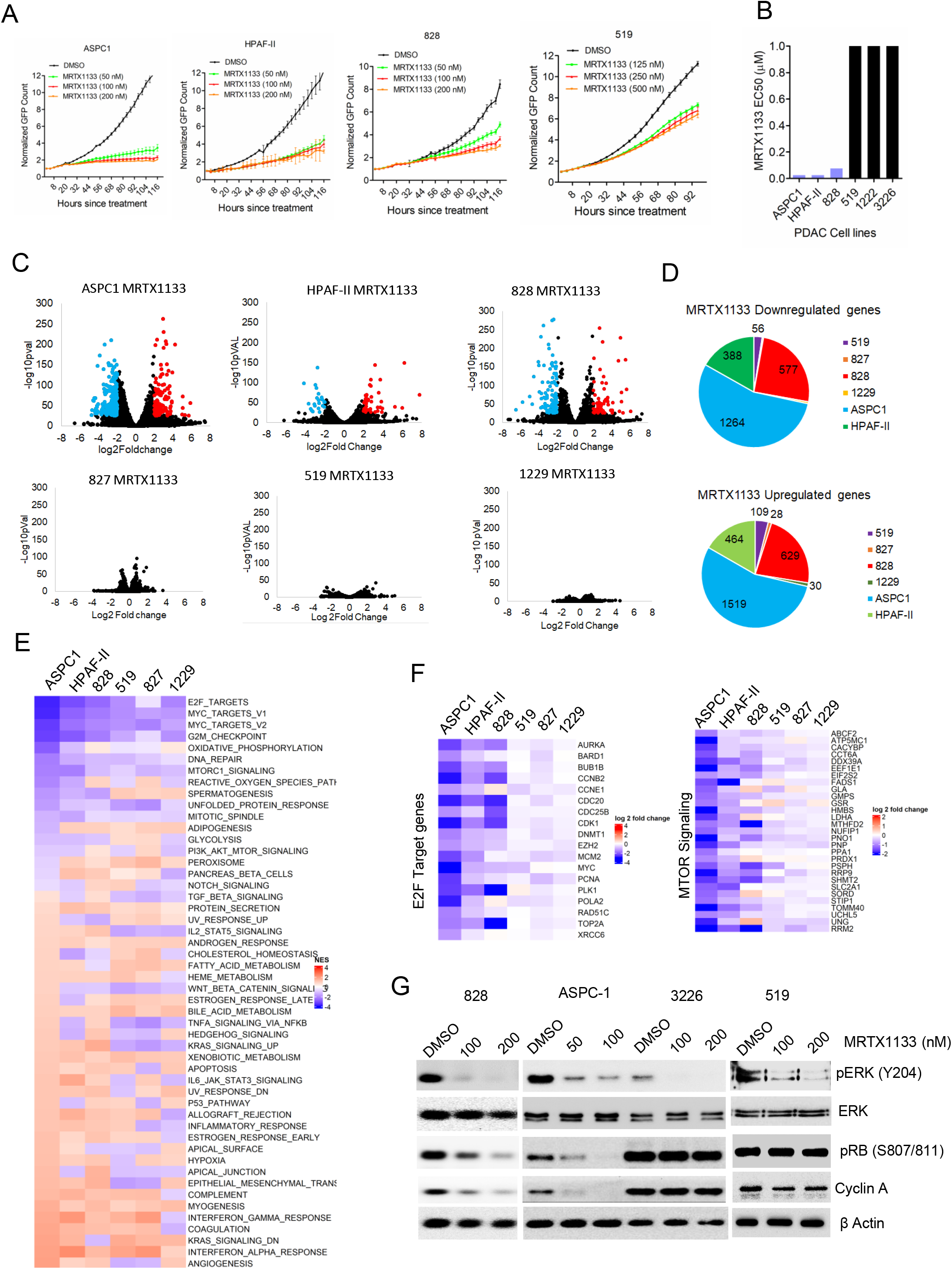
Cellular response to MRTX1133 in a panel of established and patient-derived KRAS^G12D^ mutant PDAC cell lines. (A) Live cell imaging using IncuCyte S3 on the indicated cell lines treated with different concentrations of MRTX1133. Cell proliferation was determined based on the GFP counts. Graph represents mean and SD from triplicates. Experiments were done at two independent times (B) EC50 values of MRTX1133 on different PDAC cell lines based on its impact on the cell proliferation. (C) Volcano plots indicating the differentially expressed genes on the indicated cell lines treated with MRTX1133 (100 nM) up to 48 H based on Transcriptome analysis. Blue: represents genes that are significantly downregulated (log_2_Fold change<-1.2, - log_10_pVal >2). Red represents genes that are significantly upregulated (log_2_Fold change>1.2, - log_10_pVal >2). (D) Pie charts on the indicated cell lines indicate the total number of genes that are upregulated (log_2_Fold change>1.2, -log_10_pVal >2) and downregulated (log_2_Fold change<-1.2, -log_10_pVal >2) in the presence of MRTX1133 (100 nM). (E) Heatmap depicting the GSEA hallmarks signatures that are differentially altered on different cell lines in the presence of MRTX1133 (100 nM). (F) Heatmap depicting the differential expression of the E2F target genes and the genes involved in MTOR pathway from different cell lines. (G) Immunoblotting on the indicated cell lines treated with different concentrations MRTX1133 up to 48 H.

### Genetic modifiers of the response to KRAS inhibitors

To systematically and comprehensively identify critical genes involved in resistance and sensitivity to KRAS inhibition, we employed a genome wide loss-of-function CRISPR/Cas9 library screen in a KRAS^G12D^ mutant PDAC cell line, ASPC1. Following the infection with the CRISPR/Cas9 pooled library that contains 70,948 unique single guide RNAs (sgRNAs) that target 18,053 genes with 4 sgRNAs per gene the cells were grown in the presence and absence of MRTX1133 at low concentration (6.25 nM) for 11 passages (Fig. 2A). The sgRNAs that gets depleted or enriched following the treatment with MRTX1133 imply that the loss of function of their target genes could enhance or attenuate the cytostatic effect of the drug respectively. As expected sgRNAs targeting genes that regulate ribosome biogenesis, proteasome degradation, RNA metabolism and cell cycle were significantly depleted irrespective of drug treatment following 11 passages, indicating the essential function of those genes in cell survival (Fig. S2). Moreover, a pair-wise comparison between the replicates indicated a correlation coefficient of approximately 0.9, illustrating the reproducibility of the screen (Fig. S2). Differential representation of individual sgRNA was determined based on the fold change between vehicle and MRTX1133 treated groups to define the genes that are positively and negatively selected (Fig. 2B). REACTOME pathway analyses indicated that the negatively selected sgRNAs target gene sets are highly enriched in different mitogenic pathways, including EGFR signaling, ERK signaling cascade, and metabolic pathways (Fig. 2C). We further implemented the DrugZ algorithm as previously described^38^, to calculate the NormZ values for the sgRNAs targeting each gene. Based on this analysis we identified 235 genes with NormZ values <-1.9 that were significantly depleted in the presence of MRTX1133 (Fig. 2D). The identified genes either directly regulate the function of KRAS such as SOS1 or GRB2, encompass parallel pathways to drive downstream signaling (e.g. EGFR) or act as downstream effectors such as *BRAF, MAPK3,* and *FOSL1*, suggesting that these genes contribute to resistance to MRTX1133 (Fig. 2D). The individual guides that target *EGFR* and *FOSL1*, revealed that at least 3 sgRNAs were depleted in the presence of MRTX1133 (Fig. 2E). The functional roles of EGFR and FOSL1 on the impact of MRTX1133 were further validated by gene specific depletion in ASPC1 cells. Depletion of EGFR and FOSL1 with interfering RNA significantly cooperated with MRTX1133 and yielded potent growth suppression in ASPC1 cells as monitored by live cell imaging (Fig. 2F, Fig. S2). Among the sgRNAs that were positively selected following MRTX1133 treatment, *KEAP1* was most enriched, which is consistent with previous studies (Fig. 2D) ^29, 46^. Furthermore, sgRNAs targeting *ERRFI1* and *PTEN* which act as negative regulators of EGFR singling and mTOR activity respectively were positively selected, supporting the functional importance of the mitogenic signaling pathway in bypassing KRAS inhibition (Fig. 2D). Individual guides targeting these genes further confirmed that MRTX1133 treatment increased the counts of all four sgRNAs (Fig. 2E). Moreover, since MRTX1133 renders anti-proliferative effect via RB activation, sgRNA targeting RB1 was positively selected, implying that loss of this gene could attenuate the effect of KRAS inhibition (Fig. 2D). As a validation we employed a pharmacological approach where the ASPC1 cell line was were exposed to different EGFR inhibitors in combination with MRTX1133. Based on the relative growth rate it was evident that a subset of EGFR inhibitors cooperated with MRTX1133 to induce a cytostatic response and Gefitinib significantly enhanced the efficacy of MRTX1133 in ASPC1 cells (Fig. S2). While ASPC1 is intrinsically sensitive to MRTX1133, it is expected that targeting EGFR-mediated signaling pathway could hypersensitize the cells to KRAS inhibition.

**Figure 2.**
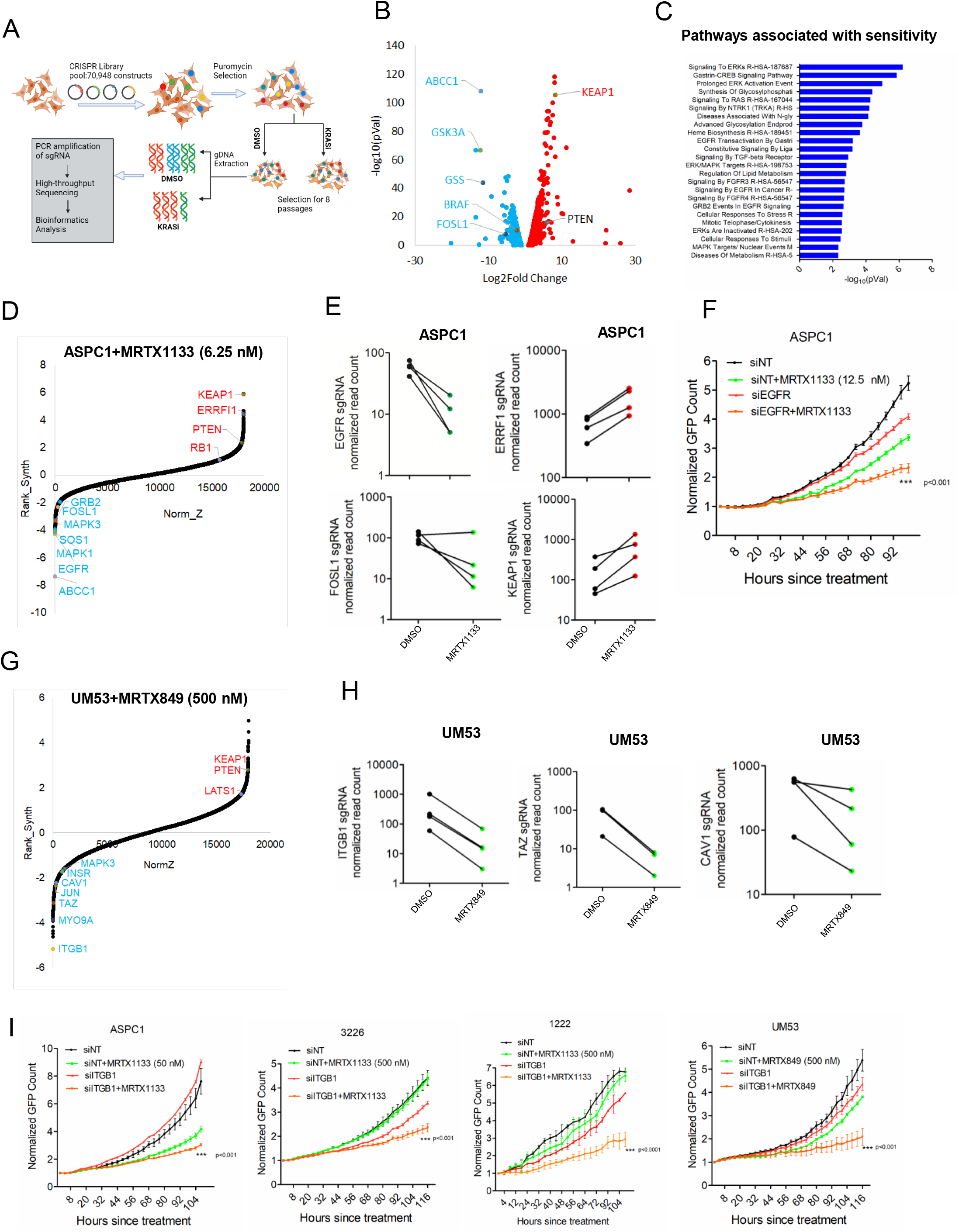
Genome wide CRISPR/CAS9 library screening in PDAC models in the presence and absence of KRAS inhibitor. (A) Schema illustrating the workflow of the CRISPR/CAS9 screening. (B) Volcano plot indicating the genes whose sgRNAs are differentially selected in the presence of MRTX1133 (6.25 nM) in ASPC1 cells. (C) ENRICHR analysis to determine the pathways that are associated with the negatively selected genes from ASPC1 cells. (D) DrugZ analysis in ASPC1 cells indicating the genes that were negatively selected during the MRTX1133 (6.25 nM) treatment. (E) Effect of MRTX1133 on the individual guides targeting, *EGFR, ERRF1, FOSL1 and KEAP1.* (F) Live cell imaging to monitor the proliferation of ASPC1 cells following EGFR deletion in the absence and presence of MRTX1133. Error bars represent mean and SD from triplicates. ***p<0.0001 as determined by two-way ANOVA. (G) DrugZ analysis in UM53 cells indicating the genes that were negatively selected during the MRTX849 (500 nM) treatment. (H) Effect of MRTX849 on the individual guides targeting, *ITGB1, TAZ and CAV1* in UM53 cells. (I) Effect of ITGB1 knockdown in ASPC1, 326, 1222 and UM53 cells in the absence and presence of MRTX1133 and MRTX849 at the indicated concentrations on cell proliferation. Graphs indicate mean SD from triplicates and the experiment was done at 2 independent times. (***p<0.0001 as determined by two-way ANOVA).

To further identify gene sets that could reverse the sensitivity to KRAS inhibition CRISPR/CAS9 screening was also performed in UM53, which is resistant to the KRAS^G12C^ inhibitor, MRTX849. Moreover, the use of this G12C model would allow us to define consistent features of KRAS inhibition with mechanistically distinct agents. Integrin subunit beta 1 (ITGB1) was identified as a negatively selected gene upon MRTX849 treatment (Fig. 2G). sgRNAS targeting, *CAV1*, TAZ and TEAD2 that are associated with integrin-mediated mechanotransdcution were also dropped out and the individual guides were depleted following MRTX849 treatment in UM53 cells (Fig. 2G & H)^47, 48^. Similarly, the individual guides against ITGB1 in ASPC1 cells were selectively depleted in the presence of MRTX1133 (Fig. S2). Further validation was performed using gene specific RNAi that target ITGB1 in combination with KRAS inhibition in 4 different PDAC cell lines. Although ASPC1 is sensitive to MRTX1133, deletion of ITGB1 modestly enhanced the efficacy of the drug (Fig. 2I). In the resistant cell lines, 3226, 1222 and UM53, ITGB1 deletion significantly reversed the sensitivity to KRAS inhibition (Fig. 2I). Long term colony formation assay further confirmed that depletion of ITGB1 enhanced the anti-proliferative effect of KRAS inhibition in 3226 and UM53 cell lines (Fig. S2). Overall our data suggest that the focal adhesion to ECM might overcome the response to KRAS inhibition. This observation is similar to a previous study, which has demonstrated that the anchorage dependent growth of KRAS mutant models can attenuate the impact of KRAS inhibition ^49^.

### Potent efficacy of MRTX1133 in 3D cell culture and in vivo settings

Since anchorage dependent cell growth could overcome KRAS dependency, we assessed the efficacy of MRTX1133 in organoids, derived from the PDAC cell lines, 3226 and 519 that are intrinsically resistant to KRAS inhibition ^13, 50^. Surprisingly, the 3226 and 519 organoids underwent a significant suppression of proliferation with MRTX1133 treatment at a concentration which had no effect in 2D culture (Fig. 3A & B). We also evaluated the effect of KRAS^G12C^ inhibitor, MRTX849, on organoids derived from UM53 cell line, which is also relatively resistant to KRAS inhibition. Similar to the G12D models, MRTX849 significantly delayed the growth of UM53 organoids (Fig. S3). Consistent with its efficacy against organoid growth, MRTX1133 displayed a cytostatic effect on tumor growth *in vivo* against two different PDX models (828 and 3226) (Fig. 3C & D). Since, the tumor microenvironment (TME) and features of the immune system play a major role in the pathogenesis of PDAC, we additionally investigated the effect of MRTX1133 in an immunocompetent setting^22^. We developed a syngeneic PDAC model derived from *Kras^+/LSL-G12D^;Pdx1-Cre* (KC) mice termed KC4568 (Fig. S4). We also employed two additional mouse syngeneic models that were previously established from KPC mouse (4662)^51^ and the iKRAS^G12D^ model where the oncogenic KRAS^G12D^ is induced by Doxycycline (AKB6)^52^. Similar to the human cell lines, the mouse models displayed differential response to MRTX1133 in cell culture. BrdU incorporation in the 4662 and KC-4568 cell lines was not inhibited in the presence of MRTX1133 indicating that these models are resistant to KRAS inhibition (Fig. 3E & S3). However, the proliferation of iKRAS AKB6 cell line in the presence of MRTX1133 was suppressed, which further resulted in the inhibition of BrdU incorporation (Fig. 3E & S3).

**Figure 3.**
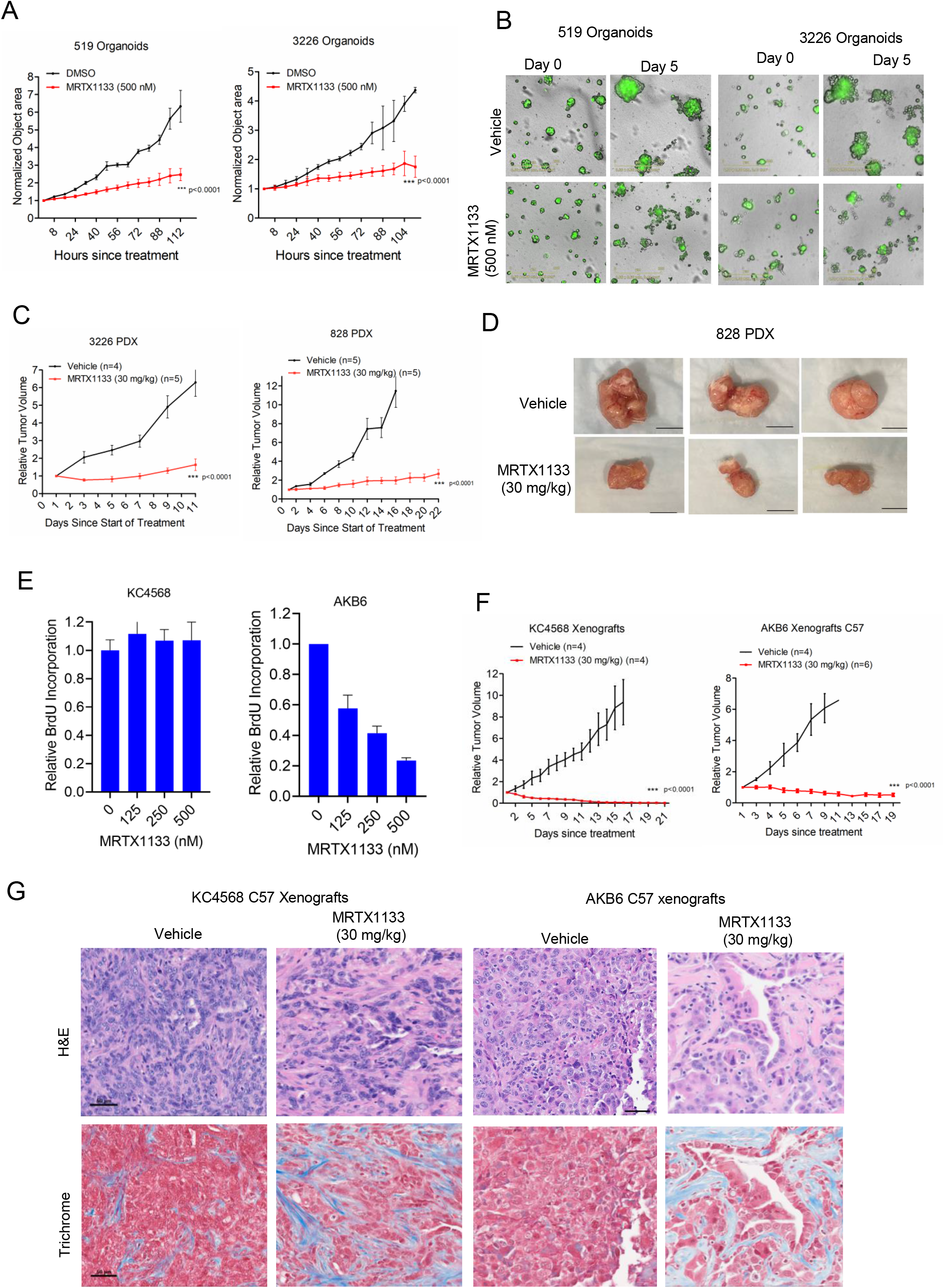
*In vivo* efficacy of MRTX1133 on PDX and mouse derived syngeneic PDAC models. (A) Effect of MRTX1133 on the growth of organoids derived from 519 PDAC cell line in the absence and presence of MRTX1133 (500 nM). Growth of organoids were determined based on the object area using IncuCyte S3. Error bars represent mean and SEM from two different experiments that were done in triplicates. (*** p<0.0001 as determined by two-way ANOVA). (B) Representative images of organoids at day 0 and day 5 in the absence and presence of MRTX1133 (500 nM). (C) Relative tumor volume of 3226 and 828 PDX following vehicle and MRTX1133 (30 mg/kg) treatment, administered intraperitonially (q.i.d) for the indicated number of days. (*** p<0.0001 as determined by two-way ANOVA). (D) Representative images of tumors excised from 828 PDX that were treated with vehicle and MRTX1133 (30 mg/kg) (E) Relative BrdU incorporation in two different syngeneic models, KC4568 and AKB6 that harbor G12D KRAS mutation following the treatment with different concentrations of MRTX1133 up to 72 H. Mean and SD were calculated from triplicates and the experiments were done at three independent times. (F) Effect of MRTX1133 (30 mg/kg) on the relative tumor growth of xenografts derived from KC4568 and AKB6 in C57/BL6 mice. (*** p<0.0001 as determined by two-way ANOVA). (G) Representative images of H&E and Masson’s Trichrome staining on tumor tissues excised from KC4568 and AKB6 xenografts that were treated with vehicle and MRTX1133 (30 mg/kg) in C57/BL6 mice.

Interestingly, similar to the human cell lines, the 4662 model possessed a more profound response to MRTX1133 when grown in organoids as compared to the 2D cell culture (Fig. S3). The *in vivo* anti-tumor effect of MRTX1133 was examined in C57BL6J mice, bearing tumors derived from the KC-4568 and AKB6 cell lines. KRAS inhibition resulted in durable disease control in both the xenografts; however, unlike the PDX models, there was significant tumor regression (Fig. 3F & S3). Furthermore, H&E staining revealed major changes in the histology with fewer tumor cell population and enhanced collagen deposition in both KC4568 and AKB6 xenografts, indicating that MRTX1133 alters TME (Fig. 3G).

### Mechanistic impact of MRTX1133 in the tumor microenvironment

To examine the molecular pathways that are perturbed by MRTX1133 in the tumor component, which could further impact the TME, we performed single cell RNA sequencing on pooled AKB6 tumor tissues that were treated with vehicle and MRTX1133. The single cell sequencing resolved multiple cell types including cancer cells, T-cells, Neutrophils, macrophages, dendritic cells, and cancer-associated fibroblasts (CAFs) within the tumors (Fig. S4 & S5). Among the different cell clusters, a particularly striking difference was selectively observed in tumor cells, T cells and neutrophils following MRTX1133 treatment (Fig. 4A, Fig. S4). The tumor population existed in two distinct subpopulations, which both express high levels of epithelial marker genes such as *Krt7, Krt8* and *Krt19.* However, in one of the subpopulations epithelial associated genes such as *Cdh1, Lamc2, Plec, Ecm1* were expressed at lower levels suggesting that a subset of tumor cells is undergoing and epithelial to mesenchymal transition (EMT) (Fig. 4B). Interestingly, both the cancer populations were dramatically decreased following MRTX1333 treatment (Fig. 4C). Further examination of differentially expressed genes in the tumor population revelated that MRTX1133 resulted in upregulation of genes such as *Cxcl9, Irgm1, Irf1* that are associated with immune-related pathways, including the activation of Interferon-γ signaling (Fig. 4D & E).

**Figure 4:**
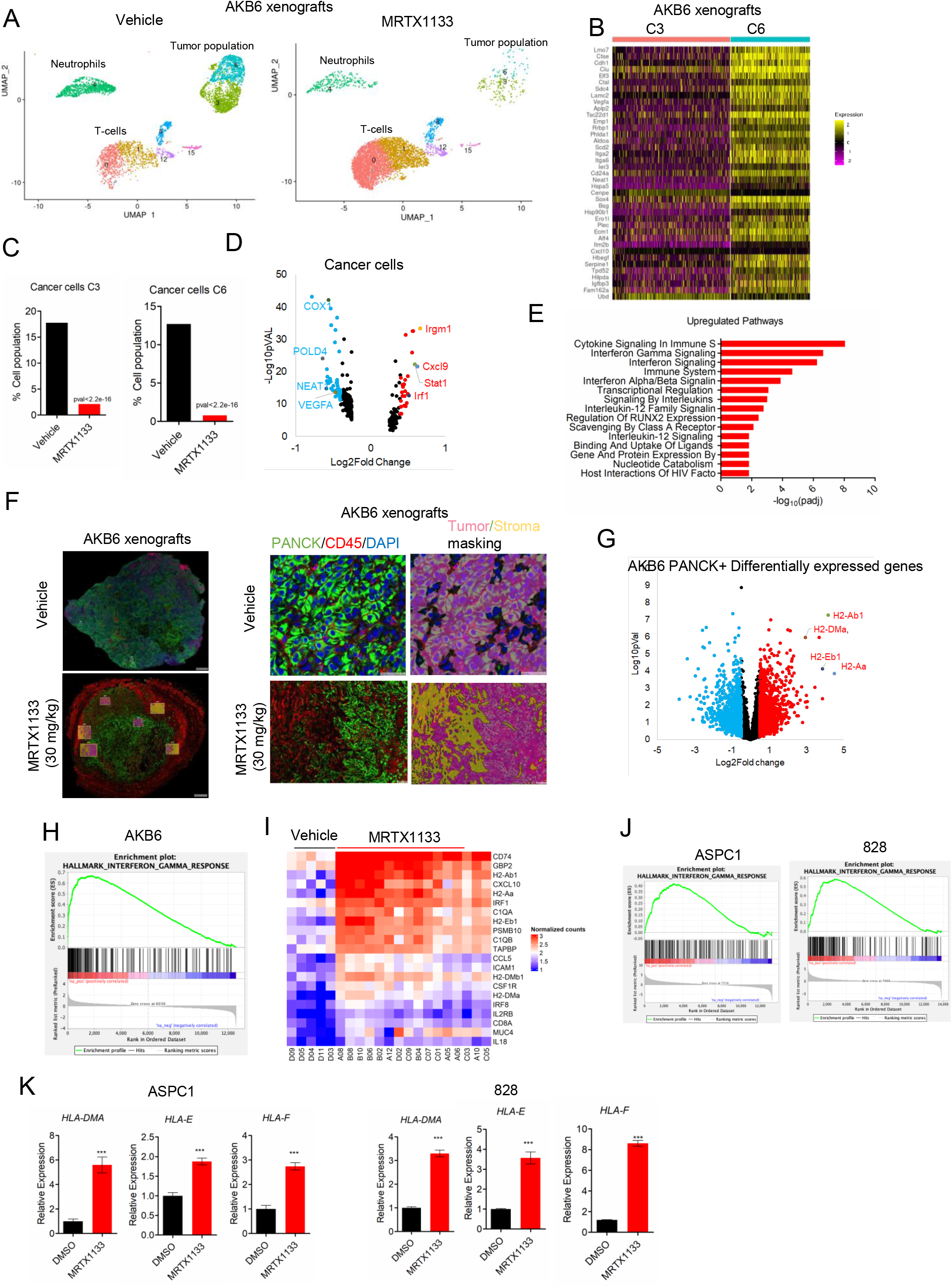
Mechanistic investigation on *in vivo* effect of MRTX1133. (A) Single-cell clustering of vehicle and MRTX1133-treated AKB6 tumors to selectively indicate Tumor cells, T-cells and neutrophils. (B) Seurat heatmap depicting the expression of genes that distinguish two different tumor population. (C) Bar graph indicating the tumor cell population from the vehicle and MRTX1133 samples. (p value was determined by Fisher extract test) (D) Volcano plot to represent the differentially expressed genes following MRTX1133 treatment from the tumor population. (E) ENRICHR analysis to identify pathways that are associated with the upregulated genes from the tumor population. (F) Representative tumor images from the tissues excised from AKB6 xenografts that were treated with Vehicle (n=2) and MRTX1133 (n=4) and stained with PACNK, CD45 and DAPI. ROIs were selected for DSP analysis. Representative image of an ROI, demarcating the tumor and stromal population based on PANCK and CD45 staining respectively and the masking was performed appropriately. (G) Volcano plots illustrate the differentially expressed genes, on the PANCK+ cells following MRTX1133 treatment on the AKB6 xenografts. Blue: represents downregulated genes (log_2_Fold change<-0.5). Red represents upregulated genes (log_2_Fold change>0.5). Antigen presenting genes were significantly upregulated and indicated. (H) GSEA analysis identified the upregulated genes following MRTX1133 treatment were significantly enriched for Interferon γ signaling. (I) Heat map depicting the upregulated genes across multiple ROIs from the vehicle treated tumor tissues (n=2) and MRTX1133 treated tumor tissues (n=4). (J) GSEA analysis based on transcriptome analysis on ASPC1 and 828 cell lines, treated with MRTX1133 (100 nM) identified that the upregulated genes were significantly enriched for Interferon γ signaling. (K) Column graph indicates the relative expression of *HLA-DMA, HLA-E* and *HLA-F* from ASPC1 and 828 cell lines treated with MRTX1133 based on transcriptome analysis. Error bar represents mean and SD. (***p<0.0001 as determine student t-test).

As a validation approach, we performed digital spatial profiling (DSP) whole transcriptome analysis on paraffin embedded tumor tissues. The tissues were stained with fluorescent tagged anti-PANCK and anti-CD45 antibodies that selectively recognizes the tumor and the stromal population respectively along with a cocktail of antibodies conjugated with DNA indexing oligonucleotides. Based on the fluorescent signal, digital masking was performed that allows to demarcate the tumor and stromal cells (Fig. 4F). Multiple regions of interest (ROIs) were selected and illuminated to release the oligos that are digitally counted and based on which the differentially expressed genes were determined (Fig. 4G). Pathway analysis on the downregulated genes against the Hallmark genesets in the Molecular Signatures Database (MSigDB)^41–44^ revealed a significant enrichment on pathways including hypoxia, glycolysis and MTOR signaling, confirming the impact of KRAS inhibition (Fig. S6). Based on the normalized counts, it was evident that MRTX1133 resulted in the suppression of the genes involved in those pathways across all the selected ROIs (Fig. S6). Similar analysis on the upregulated genes, displayed a high enrichment on different immune-related pathways, which is consistent with the single cell sequencing data (Fig. S6). GSEA analysis further confirmed that the upregulated genes were significantly enriched for Interferon-γ signaling, which resulted in the upregulation of MHC antigen presentation molecules (Fig. 4H & Fig. 4G). The critical genes involved in the interferon-like pathways such as *CD74, GBP2, H2-Ab1, CXCL10, IRF1 etc.* were prominently upregulated by MRTX1133 across all the ROIs (Fig. 4I). GSEA analysis based on the transcriptome data from ASPC1 and 828 cell lines revealed that MRTX1133-mediated upregulated genes were also significantly enriched to the Interferon-γ signaling, indicating that the impact of KRAS inhibition on the immune component is highly conserved between *in vitro* and *in vivo* settings and tumor cell autonomous (Fig. 4J & S6). The expression of *HLA-DMA, HLA-E* and *HLA-F,* which are involved in antigen presentation were significantly upregulated in ASPC1, HPAF-II and 828 cells (Fig. 4K). Together these data provide evidence that the pharmacological inhibition of KRAS upregulates the intra-tumoral antigen presentation and Interferon-like signaling pathways that are known to enhance immune infiltration to the TME^53–55^

### MRTX1133 broadly modulates the tumor immune microenvironment

We further analyzed the single cell RNA seq data to investigate the effect of MRTX1133 on other immune components in the TME. An important immune component that was modulated by MRTX1133 is the neutrophil population, which was prominently decreased in the drug treated group (Fig. 5A). Prior studies have shown that neutrophils promote tumor growth via inhibiting the activation of effector CD8^+^ T cells^56, 57^. Based on the differential gene expression in the neutrophils following MRTX1133 treatment, the downregulated genes were associated with metabolism and hypoxia (Fig. S7). Expression of genes involved in these pathways such as *Hilpda, Ldha, Eno1* were significantly downregulated by MRTX1133, which could inhibit the survival of neutrophils and enhance T-cell activation (Fig. 5B & C). Additionally, the increase in antigen presentation in tumor cells could also recruit T cells. Consistent to this notion, MRTX1133 treatment notably increased the T-cell populations that comprise, CD4^+^/ CD8^+^ T cells and natural killer T cells (NKT) (Fig. 5D & Fig. S4). Among the different T-cell population, a prominent increase was observed in the CD8^+^ T cells (Fig. 5E). Increase in interferon gama (*Ifng)* was selectively observed in the CD8^+^ T-cells following MRTX1133 treatment and the upregulation of *Icos and CD69* was observed in both CD4^+^ /CD8^+^ T cells and NKT cells, indicating the activation of effector T cells to induce anti-tumor immunity (Fig. 5D & 5F, Fig. S7)^22^. Moreover, the cytolytic function of T-cells was enhanced by MRTX1133, as evident by the upregulation of *Prf1* in the CD8^+^ T cells and *Klrb1b* in the NKT cells (Fig. 5D, Fig. S7). To validate this observation, tumor tissues from the vehicle and MRTX1133 treated groups were subjected to multi-spectral staining that comprises Pan-CK, DAPI and the T-cell marker, CD8. Consistent to the single cell sequencing data, MRTX1133-treated tissues exhibited a significantly higher CD8 expression in both the tumor and stroma confirming that MRTX1133 induces T-cell activation (Fig. 5G). CAFs and macrophages, which are also an important immune component in the TME were not notably modulated by MRTX1133 (Fig. S7). Gene expression of markers specific for iCAFS (*C1s1, Col14a1, Has1)* and myCAFs (*Col12a1)* were not altered following MRTX1133 treatment (Fig. S7)^22^. We also evaluated the effect of MRTX1133 on AKB6 xenografts developed in immune deficient NSG mice. Although MRTX1133 induced a cytostatic response against tumor growth, the disease control was more profound in immune-competent mice (Fig. 5H). Overall our data suggests that oncogenic KRAS inhibition by MRTX1133 possess a major impact on TME and enhance cytotoxic T-cell infiltration to induce a durable disease control.

**Figure 5:**
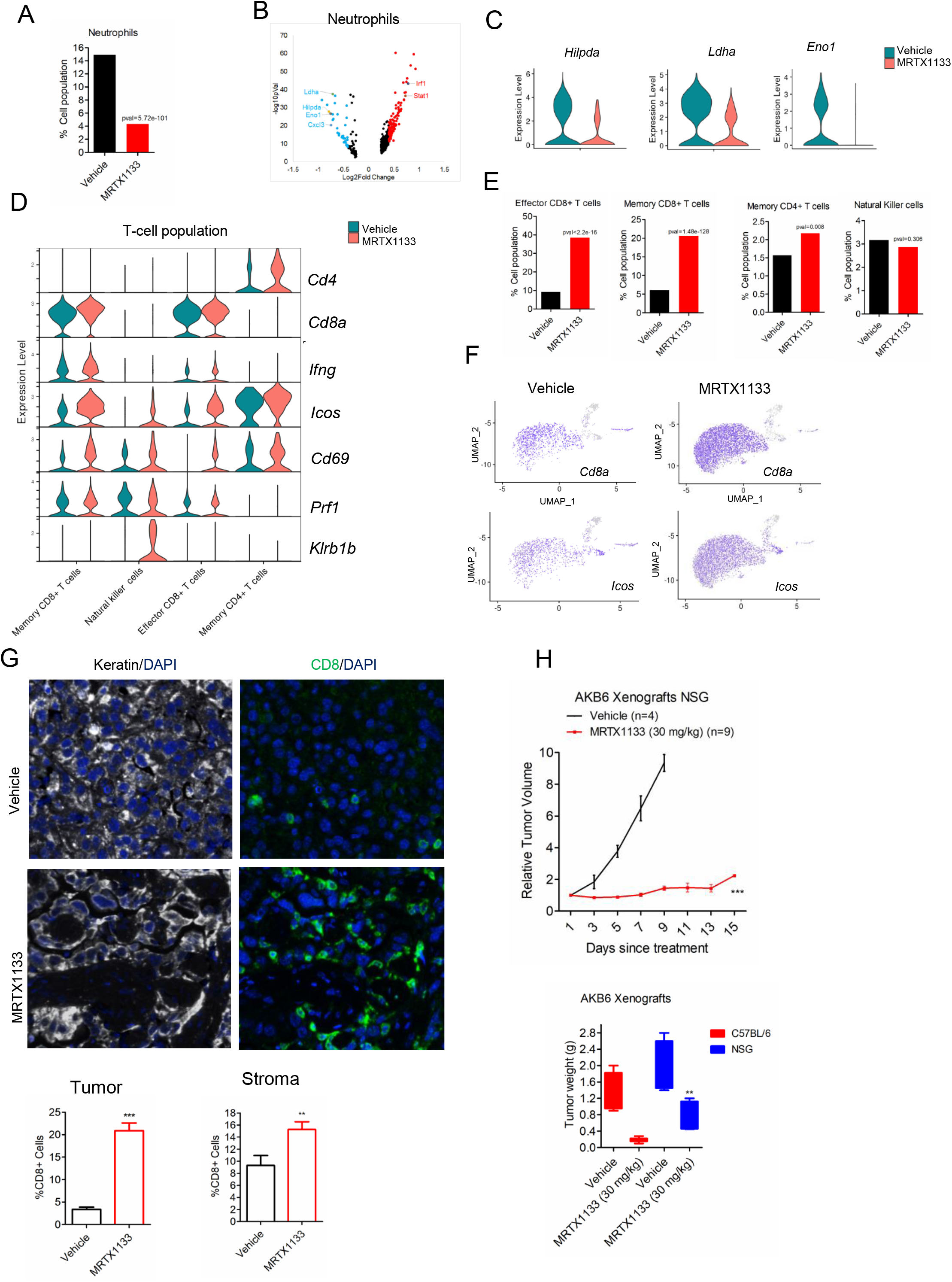
Single cell sequencing analysis on the immune component from AKB6 xenografts. (A) Column graph indicating the neutrophils population in the absence and presence of MRTX1133. (p value was determine by Fisher extract test). (B) Volcano plot indicating the downregulated and upregulated genes on the neutrophils population following MRTX1133 treatment. (C) Violin plots indicating the expression of indicated genes within the neutrophil population in the presence and absence of MRTX1133. (D) Violin plots to indicate the differential expression of T-cell markers between the vehicle and MRTX1133-treated groups. (E) Column graphs indicating the different T-cell population that comprise Memory CD8^+^ T cells, Effector CD8^+^ T cells, NKT cells and Memory CD4^+^ T cells following MRTX1133 treatment. P values were determined by Fisher extract test. (F) Seurat feature plots to illustrate the differential expression of representative T-cell markers from the vehicle and MRTX1133-treated groups. (G) Representative images from the multispectral staining on the AKB6 tumors (PanKeratin, CD8 and DAPI). Column graph represent the fraction of CD8-positive cells from the stroma and tumor following the treatment with MRTX1133. Mean and SEM were determined from 6 ROIs from vehicle (n=3) and MRTX1133 (n=3) treated tumor tissues. (***p<0.0001 and **p<0.01 as determine by student t-test). (H) *In vivo* effect of MRTX1133 on the tumor growth of AKB6 xenografts in NSG mice. Tumor weights from the AKB6 xenografts derived from NSG and C57/BL6 strains treated with vehicle and MRTX1133. Mean and SEM were shown. (*** p<0001 as determined by student t test).

## Discussion

The mutation of KRAS is considered the initiating oncogenic driver for the pathogenesis of PDAC^9, 58^. While multiple different KRAS mutant isoforms occur in PDAC, the G12D mutation is the most common occurring ∼40% of cases^29, 59^. Multiple targeted therapies have been interrogated that inhibit KRAS-mediated downstream effector pathways in clinical trials, although the therapeutic benefits are limited^60, 61^. While KRAS was considered undruggable for many years, structure-based drug design approaches have led to the development of multiple selective pharmacological inhibitors ^28, 62^.

The current work demonstrates the cellular response of a potent KRAS^G12D^ inhibitor, MRTX1133 in different patient-derived PDAC cell lines and syngeneic models. The efficacy of MRTX1133 *in vitro* cell culture assays is heterogeneous among the PDAC models and characterized as extremely sensitive or largely totally refractory. Based on our transcriptome analysis, the top pathway that is significantly inhibited in the sensitive models is the cell cycle machinery, which remained largely unperturbed in the resistant models, suggesting the involvement of parallel pathways that allow to bypass the impact of KRAS inhibition and drive cell cycle. Consistent with this concept key mediators of KRAS and RTK signaling were identified as cooperating with pharmacological mutant KRAS inhibitors using CRISPR/Cas9 screening. The genes identified included those directly associated with controlling RAS activity that could represent activity through other RAS genes, as well as downstream effectors (e.g. BRAF and MAPK3). Conversely, EGFR was a key mediator of resistance that could be further targeted pharmacologically to block the ability of cells to bypass mutant KRAS inhibition. Genes whose loss associated with resistance are known to further deregulate signaling pathways (e.g. PTEN) or limit effects on cell cycle (e.g. RB). These findings advance a number of combinatorial strategies that could be deployed with MRTX1133 to enhance cell autonomous activity on tumor cell division.

Our CRISPR screen analysis further revealed a cadre of genes whose depletion potently cooperated with KRAS inhibition (MRTX1133 and MRTX849) that are associated with adhesion mediated signaling. This includes integrins (ITGB1 and ITGAV) and genes involved in mechanotransdcution (CAV1, YAP, TEAD2)^47, 48, 63^. Our study further illustrates that the resistance to MRTX1133 in 2D culture could be alleviated when the cells are grown in organoids, and this phenomenon was observed in both human and mouse PDAC models. Presumably, integrin-mediated mechano-transduction is limited in the Matrigel matrix, thus growth of organoids is considerably inhibited following KRAS inhibition. It has been posited that cell culture yields enhanced sensitivity to therapeutic agents while the organoids or xenografts are considered a more stringent system to assess clinically relevant drug efficacy^64^. However, in the context of KRAS inhibition, cells cultured as organoids are considerably more sensitive to therapeutic targeting^13, 49^. The different PDAC models were also evaluated as xenografts and a prominent anti-tumor effect was observed following the treatment with MRTX1133. Interestingly, prior studies using different targeted agents or genetic depletion of KRAS did not indicate a drastic difference in efficacy between the 2D cell culture and organoids ^32, 65^. However, the data here highlight the importance of the 3D cell culture or organoids to be used in an *in vitro* setting to more accurately predict the efficacy of KRAS inhibitors against tumor growth.

Although MRTX1333 inhibited tumor growth in all the G12D mutant models tested, a significant difference on disease control was observed between immune-deficient NSG strain and an immune-competent C57BL/6 strain. While loss of KRAS was known to impact the TME, our study illustrates that pharmacological inhibition of KRAS could yield a similar effect^22^. Single cell sequencing revealed highly significant changes in TME following the treatment with MRTX1133. Suppression of tumor cell population along with increase in cytolytic and effector CD8^+^/CD^4+^ T cells and decrease in neutrophils implies that MRTX1133-mediated tumor regression is associated with TME remodeling. In this study we elucidated the mechanism through which MRTX1133 enhances the infiltration of effector T cells by utilizing a more sophisticated unbiased approach, Digital spatial Profiling (DSP) to determine the differential gene expression selectively on the tumor population. This technique is more advanced than the conventional bulk RNA sequencing, because it spatially discriminates the tumor population from the stromal cells^66^. Based on our analysis from both single cell RNA sequencing and DSP, the tumor cells displayed a prominent upregulation of genes involved in interferon γ signaling and MHC class II antigen presentation. These molecular events have been proposed to enhance T-cell recognition, activation of effector CD8^+^ T cells and infiltration to eliminate the cancer cells^53, 67–69^. Neutrophils, one of the major components in the TME are known to promote tumorigenesis by suppressing the function of effector T-cells^57^. The survival of neutrophils is mediated by oxidative mitochondrial metabolism that results in a hypoxic condition and enhances Reactive Oxygen Species (ROS) production to inhibit effector T-cell activation and suppress the anti-tumor immunity^56^. The impact of MRTX1133 on tumor cells downregulates genes associated with hypoxia and metabolism in neutrophil population that impacts their viability and promotes T-cell activation.

The impact on TME and activation of effector T cells are direct consequences of KRAS inhibition, because in our prior studies the treatment with a MEK inhibitor in combination with CDK4/6 inhibitor did not result in tumor regression rather the tumor escaped the acute cytostatic effect ^53^. The MRTX1133-mediated tumor regression is reversible because cessation of treatment caused tumor recurrence in one of the syngeneic models (data not shown), which is consistent with a recent study^59^. Moreover, our single cell sequencing data revealed an increase in the exhaustion markers such as *Ctla4* and *Lag3* that suppress the anti-tumor immunity, suggesting new combinatorial treatment modalities involving concurrent targeting of KRAS and immunotherapy (Fig. S7).

In summary, this study highlights the efficacy of a KRAS^G12D^ inhibitor, MRTX1133 in PDAC models and demonstrates a differential response in 2D cell culture, organoids and *in vivo.* While we demonstrate two distinct molecular components, focal adhesion and TME remodeling that modulate the response to MRTX1133, it is limited to preclinical setting. Whether the genes involved in those pathways could serve as biomarkers of response or resistance to KRAS inhibitors in patient models need further investigation.

## Methods

### Cell Culture and Therapeutic agents

Human primary PDAC cell lines 828, 519, 3226, 1222, and 827 and mouse primary PDAC cell line, KC4568 were grown in Keratinocyte Serum Free medium (KSF), supplemented with 2% FBS, EGF (0.2 ng/mL) and bovine pituitary extract (30 µg/mL) (Life Technologies, Carlsbad, CA) as described before^30^. The TC dishes were precoated with collagen (Millipore, Burlington, MA). UM53, ASPC1 and HPAF-II cell lines were kindly provided by Dr. Ethan Abel (Roswell Park Cancer Center) and were grown in RPMI medium, supplemented with 10% FBS. Mia PaCa-2 cell line was cultured in DMEM media, supplemented with 10% FBS. Mouse KPC PDAC cell line, 4662 was obtained from Dr. Vonderheide’s laboratory (University of Pennsylvania) and grown in RPMI media, supplemented with 10% FBS. The iKRAS AKB6 was kindly provided by Dr. Prasenjit Dey (Roswell Park Cancer Center) and cultured in RPMI medium, supplemented with 10% FBS. All cell lines were maintained in 37°C with 5% CO_2_ and tested to be Mycoplasma free. ASPC1, Mia PaCa-2 and HPAF-II were authenticated by STR profiling. MRTX1133 was purchased from Chemietek (Cat # CT-MRTX1133) and dissolved in DMSO to get a final concentration of 10 mM. For *in vivo* work, MRTX1133 was kindly provided by Mirati Therapeutics. Gefitinib was purchased from SelleckChem.

### Cell Proliferation Assay

To monitor the cell growth, we employed live cell imaging systems, IncuCyte S3 and CellCyte X. The cell lines were transfected to stably express H2B-GFP and seeded in 96 well TC dish in the presence and absence of test agents. The GFP counts were measured in real time that corresponds to proliferation rate.

### Knockdown experiments

The cell lines were reverse transfected with gene specific RNAis that target *ITGB1, FOSL1 and EGFR* as described in previous study^31^. On-target plus human RNAi for *ITGB1* was purchased from Horizon Discovery (Cat # L-004506-00-0005). siRNAs for *EGFR* (Cat # S563) and *FOSL1* (Cat # S15583) were purchased from Thermo Scientific. Following transfection, the cells were treated with MRTX1133 and the cell proliferation was monitored using live cell imaging.

### Western Blotting

Whole cell extracts were prepared as described in our previous studies ^30, 32^. The primary antibodies for pERK and pRB (S807/811) (Cat # 8516S) were purchased from Cell Signaling. β Actin antibody was purchased from R&D Systems (Cat # MAB8929). Cyclin A antibody was purchased from Sigma (Cat # C4710).

### Colony Formation assay

3226 and UM53 cells were seeded in 6 well dish (1000 cells/well) and treated with MRTX1133 and MRX849 respectively in combination with ITGB1 RNAi. Colonies were allowed to form for 10 days and stained with Crystal Violet.

### Organoid Cell culture

The H2B-GFP labelled 519, 3226 and UM53 cells were seeded in 50 % Matrigel basement layer (4000 cells/well) and allowed to form Organoids. Following the treatment with MRTX1133 and MRTX849, the growth of organoids were monitored using live cell imaging that determined the GFP area.

### Mice and patient-derived xenografts

NSG and C57/BL6 were maintained at Roswell Park Cancer Center animal care facilities. All animal care, drug treatments and sacrifice were approved by the Roswell Park Cancer Center Institutional Animal Care and Use Committee (IACUC) in accordance with the NIH guide for the care and use of laboratory animals. Mice were subcutaneously implanted with early passage 828 and 3226 PDX tumor fragments. AKB6 xenografts were developed in both NSG and C57/BL6 strains by subcutaneously injecting 5X10^6 cells/mouse. The entire experimental cohort was comprised of both male and female mice. The mice were always maintained in Doxycycline water to induce the oncogenic KRAS. KC4568 xenografts were developed in the C57/BL6 strain by subcutaneously injecting 5X10^6 cells/mouse. Once the tumor reached 200 mm^3^, mice were randomized in a non-blinded manner, comprising, Vehicle (n=5), and MRTX1133 (30 mg/kg) (n-5), administered intraperitoneally once per day for three weeks. MRTX1133 was dissolved in 50 mM citrate buffer and 10% Captisol. Tumor growth was monitored every other day using digital calipers and the tumor volume was calculated using the formula (length*widith*width)/2 as described before ^30^. Mice were sacrificed and the tumors were embedded in paraffin for further analysis. Mice body weights were monitored regularly to evaluate any adverse effects. Mice that died during the course of treatment were excluded from analysis. The animal

### Transcriptome Analysis

RNA was extracted using Quiagen RNeasyplus kit and analyzed via the RNA6000 Nano assay, and with the Agilent 2200 TapeStation (Agilent, CA, USA) for determination of an RNA Integrity Number (RIN), and only the cases with RIN > 7.0 were included in this study. cDNA synthesis was performed using random hexamers to obtain full-length, strand-specific representation of nonribosomal RNA transcripts. Targeted RNA sequencing libraries were prepared with the DriverMap Human Genome-Wide Gene Expression Profiling Sample Prep Kit hDM18Kv3 (Cellecta Inc., CA, USA). A defining feature of the DriverMap method is the application of pre-designed multiplex PCR primer sets targeted to specific 50-75 bp regions of all known protein-coding genes. Notably, each target-specific primer consists of a complementary sequence to specific mRNA targets plus a universal primer binding site (anchor). Ligation of oligonucleotides via PCR amplification introduces adaptors required for sequencing and sample-specific "barcodes" that flank the target sequence and are inserted into standard Illumina adaptors to permit dual-index sequencing and deconvolution of sample-specific reads using standard Illumina software.

Briefly, to mitigate primer dimer formation, anchor PCR was performed with an initial hot start at 95°C for 5 min, followed by 15 cycles of (95°C – 0.5 min, 68°C – 1 min, 72°C – 1 min), and ended with a final 10 min extension at 72°C. The reaction products were confirmed on an agarose gel in triplicate to assess replicability. PCR products were then purified by SPRI (Agentcourt, 1:1 sample: reagent ratio) and quantified with the Qubit fluorescence assay (Qubit dsDNA HS Assay Kit, ThermoFisher Scientific, MA, USA). Target-enriched RNA-seq libraries were analyzed on an Illumina NextSeq 500 sequencer using a NextSeq500/550 High Output v2 Kit (75 cycles) according to the standard manufacturer’s protocol (Illumina, CA, USA).

For RNA-seq, alignment was performed using STAR v2.7.10b, and this pipeline outputs gene-level read counts directly^33^. Transcript abundance estimates for each sample were performed using Salmon, an expectation-maximization algorithm using the UCSC gene definitions. Raw read counts for all samples were normalized using the "weighted" Trimmed Mean of M-values (TMM) approach in the Bioconductor package EdgeR^34^. After trimming the data [5% for the A values, log ratio 0.3 for the M values to a reference array (the library whose upper quartile is closest to the mean upper quartile)], scaling factors for each sample were generated using the calcNormFactors function. Genes with an average of less than ten read counts across all samples were excluded from further analysis. Following data integration, systematic bias was corrected using ComBat, as described previously^35^. DE gene (DEG) analysis was performed with EdgeR and selected based on p-value and log2 fold change (log2FC), respectively.

### CRISPR screening

Human Toronto KnockOut (TKO) CRISPR library-version 3, containing 70,948 unique sgRNAs, which targets 18,053 genes was packaged in lentivirus according to the previously published protocol^36^. ASPC1 and UM53 cells were infected with the TKOv3 lentiviral particles at an MOI ∼0.3 and the positive clones were selected using puromycin to achieve a mutant pool comprising at least 200-fold coverage of the library. The resulting mutant pool was grown in the absence and presence of KRASi at the EC25 concentration for at least 8 passages. Genomic DNA was extracted and subjected to sequencing library construction by amplifying the gRNA inserts using a 2-Step PCR as described in the previously published study^36^. Resulting libraries were further sequenced on an Illumina’s NexSeq platform. Fastq files were first subjected to adapter removal using Trim Galore (v0.6.7, https://github.com/FelixKrueger/TrimGalore). Next, the MAGeCK^37^ pipeline was used to process the adapter trimmed fastq files and the resultant count files are used as input to DrugZ^38^ python script for chemogenetic interactions identification. An in-house developed R script was used to creating plots for results visualization.

### 10X Single cell RNA sequencing

Tumors excised from AKB6 xenografts that were treated with vehicle (n=2) and MRTX1133 (n=2) were digested using Liberase (Sigma Cat # 05401020001) to get single cells and sent to RPCI’s Genomics Shared Resources for sequencing using 10X Chromium and Illumina’s NovaSeq instruments. The raw fastq data was processed using 10X’s cellranger pipeline (v6.1.2). The resultant ‘filtered_feature_bc_matrix.h5’ files for treatment and vehicle control, respectively, were used as inputs for a custom R (v4.2.0) script which utilizes the Seruat^39^ R package (v4.2.0) from CRAN. We filtered each dataset with the following criteria: percent.mt < 10, nFeature_RNA between 200 and 4000, percent.largest.gene < 20. With these parameters, we obtained 9307 and 7204 single cells in the treated and control samples, respectively. Since the treated sample resulted more single cells, we down-sampled it by randomly selecting the same number of cells (7204) as in the control. The two datasets were then integrated by using the FindIntegrationAnchors and IntegrateData function from the Seurat suite. After this step, the standard Seurat workflow was followed to cluster the combined dataset. We identified 20 clusters and annotated the clusters with ScType^40^

### Digital Spatial Profiling (DSP)

We used NanoStrings GeoMx instrument and the Mouse NGS Whole Transcriptome Atlas RNA (version 1.0) kit for ROIs selection and mRNA library preparation. We used PanCK and CD45 antibodies cocktail for staining of cancer cell and stromal segments respectively. The resulting mRNA samples are subjected to sequencing using Illumina’s NavoSeq instrument. Raw fastq files along with the configuration file are used as input to the geomxngspipeline command line tool running on a Linux server. We used NanoString’s GeomxTools Bioconductor package (https://www.bioconductor.org/packages/release/bioc/html/GeomxTools.html) for downstream data analyses. These include QC & pre-processing, data normalization, unsupervised clustering and differential gene expression. For QC, we used values for these parameters: minSegmentReads = 1000, percentTrimmed = 80, percentStitched = 80, percentAligned = 80, percentSaturation = 50, minNegativeCount = 2, maxNTCCount = 1000, minNuclei = 100, minArea = 5000, minLOQ = 2. For segments passed these QC parameters, we utilized Q3 normalization and mixed effect modeling for identifying differentially expressed genes in the PanCK+ segments between treated and control.

### Pathway analysis

We used the Enrichr^41–43^ webserver and MSigDB^44^ Hallmark 2020 dataset for all pathway analysis in this paper.

### Multi-spectral immunofluorescence (mIF) staining

Formalin-fixed Paraffin-embedded (FFPE) 4 µm sections were cut and placed on charged slides. The multispectral immunofluorescent (mIF) staining process involved serial repetitions of the following for each biomarker: epitope retrieval/stripping with ER1 (citrate buffer pH 6, AR996, Leica Biosystems) or ER2 (Tris-EDTA buffer pH9, AR9640, Leica Biosystems), blocking buffer (AKOYA Biosciences,), primary antibody, Opal Polymer HRP secondary antibody (AKOYA Biosciences), Opal Fluorophore (AKOYA Biosciences). All AKOYA reagents used for mIF staining come as a kit (NEL821001KT). Spectral DAPI (AKOYA Biosciences) was applied once slides were removed from the BOND. They were cover slipped using an aqueous method and Diamond antifade mounting medium (Invitrogen ThermoFisher). The mIF panel consisted of the following antibodies, CD8 (EPR20305, Abcam, 570), Pan Keratin (Wide Spectrum cytokeratin, Abcam, 480), and DAPI. Slides were imaged on the PhenoImager HT^®^ Automated Quantitative Pathology Imaging System (AKOYA Biosciences). Further analysis of the slides was performed using inForm^®^ Software v2.6.0 (AKOYA Biosciences).

## Acknowledgements

The author thank all members of the laboratory group and colleagues in the discussion and preparation of the manuscript. This work was supported by grants from the National Institutes of Health, National Cancer Institute (CA267467 and CA211878).

## Author contributions

Study concept and design: VK, CF, EA, PD, ESK and AKW Acquisition of data: VK, CF, JW, YW, AD and HR

Analysis and interpretation of data: VK, CF, JW, YW, AD, HR, ESK and AKW Study supervision: ESK and AKW.

## References

1 Yao, W., Maitra, A. & Ying, H. Recent insights into the biology of pancreatic cancer. EBioMedicine 53, 102655, doi:10.1016/j.ebiom.2020.102655 (2020).

2 Kleeff, J. et al. Pancreatic cancer. Nat Rev Dis Primers 2, 16022, doi:10.1038/nrdp.2016.22 (2016).

3 Neoptolemos, J. P. et al. Therapeutic developments in pancreatic cancer: current and future perspectives. Nat Rev Gastroenterol Hepatol 15, 333–348, doi:10.1038/s41575-018-0005-x (2018).

4 Stott, M. C., Oldfield, L., Hale, J., Costello, E. & Halloran, C. M. Recent advances in understanding pancreatic cancer. Fac Rev 11, 9, doi:10.12703/r/11-9 (2022).

5 Witkiewicz, A. K. et al. Whole Exome Sequencing of Pancreatic Cancer:Genetic Diversity, Prognostic Features, and Potential Therapeutic Targets. Nature Communication (2014).

6 Bailey, P. et al. Genomic analyses identify molecular subtypes of pancreatic cancer. Nature 531, 47–52, doi:10.1038/nature16965 (2016).

7 Knudsen, E. S., O’Reilly, E. M., Brody, J. R. & Witkiewicz, A. K. Genetic Diversity of Pancreatic Ductal Adenocarcinoma and Opportunities for Precision Medicine. Gastroenterology, doi:10.1053/j.gastro.2015.08.056 (2015).

8 Infante, J. R. et al. A randomised, double-blind, placebo-controlled trial of trametinib, an oral MEK inhibitor, in combination with gemcitabine for patients with untreated metastatic adenocarcinoma of the pancreas. Eur J Cancer 50, 2072–2081, doi:10.1016/j.ejca.2014.04.024 (2014).

9 Huang, L., Guo, Z., Wang, F. & Fu, L. KRAS mutation: from undruggable to druggable in cancer. Signal Transduct Target Ther 6, 386, doi:10.1038/s41392-021-00780-4 (2021).

10 Waters, A. M. & Der, C. J. KRAS: The Critical Driver and Therapeutic Target for Pancreatic Cancer. Cold Spring Harb Perspect Med 8, doi:10.1101/cshperspect.a031435 (2018).

11 Nollmann, F. I. & Ruess, D. A. Targeting Mutant KRAS in Pancreatic Cancer: Futile or Promising? Biomedicines 8, doi:10.3390/biomedicines8080281 (2020).

12 Zeitouni, D., Pylayeva-Gupta, Y., Der, C. J. & Bryant, K. L. KRAS Mutant Pancreatic Cancer: No Lone Path to an Effective Treatment. Cancers (Basel) 8, doi:10.3390/cancers8040045 (2016).

13 Muzumdar, M. D. et al. Survival of pancreatic cancer cells lacking KRAS function. Nat Commun 8, 1090, doi:10.1038/s41467-017-00942-5 (2017).

14 Kapoor, A. et al. Yap1 activation enables bypass of oncogenic Kras addiction in pancreatic cancer. Cell 158, 185–197, doi:10.1016/j.cell.2014.06.003 (2014).

15 Viale, A. et al. Oncogene ablation-resistant pancreatic cancer cells depend on mitochondrial function. Nature 514, 628–632, doi:10.1038/nature13611 (2014).

16 Collins, M. A. et al. Oncogenic Kras is required for both the initiation and maintenance of pancreatic cancer in mice. J Clin Invest 122, 639–653, doi:10.1172/JCI59227 (2012).

17 Shinkawa, T., Ohuchida, K. & Nakamura, M. Heterogeneity of Cancer-Associated Fibroblasts and the Tumor Immune Microenvironment in Pancreatic Cancer. Cancers (Basel) 14, doi:10.3390/cancers14163994 (2022).

18 Hosein, A. N., Brekken, R. A. & Maitra, A. Pancreatic cancer stroma: an update on therapeutic targeting strategies. Nat Rev Gastroenterol Hepatol 17, 487–505, doi:10.1038/s41575-020-0300-1 (2020).

19 Rhim, A. D. et al. Stromal elements act to restrain, rather than support, pancreatic ductal adenocarcinoma. Cancer Cell 25, 735–747, doi:10.1016/j.ccr.2014.04.021 (2014).

20 Olive, K. P. et al. Inhibition of Hedgehog signaling enhances delivery of chemotherapy in a mouse model of pancreatic cancer. Science 324, 1457–1461, doi:1171362 [pii] 10.1126/science.1171362 (2009).

21 Li, X. et al. Immune checkpoint blockade in pancreatic cancer: Trudging through the immune desert. Seminars in cancer biology 86, 14–27, doi:10.1016/j.semcancer.2022.08.009 (2022).

22 Ischenko, I. et al. KRAS drives immune evasion in a genetic model of pancreatic cancer. Nat Commun 12, 1482, doi:10.1038/s41467-021-21736-w (2021).

23 Nakajima, E. C. et al. FDA Approval Summary: Sotorasib for KRAS G12C-Mutated Metastatic NSCLC. Clin Cancer Res 28, 1482–1486, doi:10.1158/1078-0432.CCR-21-3074 (2022).

24 Strickler, J. H. et al. Sotorasib in KRAS p.G12C-Mutated Advanced Pancreatic Cancer. The New England journal of medicine 388, 33–43, doi:10.1056/NEJMoa2208470 (2023).

25 Ning, W., Marti, T. M., Dorn, P. & Peng, R. W. Non-genetic adaptive resistance to KRAS(G12C) inhibition: EMT is not the only culprit. Frontiers in oncology 12, 1004669, doi:10.3389/fonc.2022.1004669 (2022).

26 Xue, J. Y. et al. Rapid non-uniform adaptation to conformation-specific KRAS(G12C) inhibition. Nature 577, 421–425, doi:10.1038/s41586-019-1884-x (2020).

27 Witkiewicz, A. K. et al. Whole-exome sequencing of pancreatic cancer defines genetic diversity and therapeutic targets. Nat Commun 6, 6744, doi:10.1038/ncomms7744 (2015).

28 Wang, X. et al. Identification of MRTX1133, a Noncovalent, Potent, and Selective KRAS(G12D) Inhibitor. Journal of medicinal chemistry 65, 3123–3133, doi:10.1021/acs.jmedchem.1c01688 (2022).

29 Hallin, J. et al. Anti-tumor efficacy of a potent and selective non-covalent KRAS(G12D) inhibitor. Nat Med 28, 2171–2182, doi:10.1038/s41591-022-02007-7 (2022).

30 Knudsen, E. S. et al. Cell cycle plasticity driven by MTOR signaling: integral resistance to CDK4/6 inhibition in patient-derived models of pancreatic cancer. Oncogene, doi:10.1038/s41388-018-0650-0 (2019).

31 Knudsen, E. S. et al. CDK/cyclin dependencies define extreme cancer cell-cycle heterogeneity and collateral vulnerabilities. Cell Rep 38, 110448, doi:10.1016/j.celrep.2022.110448 (2022).

32 Kumarasamy, V., Ruiz, A., Nambiar, R., Witkiewicz, A. K. & Knudsen, E. S. Chemotherapy impacts on the cellular response to CDK4/6 inhibition: distinct mechanisms of interaction and efficacy in models of pancreatic cancer. Oncogene 39, 1831–1845, doi:10.1038/s41388-019-1102-1 (2020).

33 Dobin, A. et al. STAR: ultrafast universal RNA-seq aligner. Bioinformatics 29, 15–21, doi:10.1093/bioinformatics/bts635 (2013).

34 Robinson, M. D. & Oshlack, A. A scaling normalization method for differential expression analysis of RNA-seq data. Genome Biol 11, R25, doi:10.1186/gb-2010-11-3-r25 (2010).

35 Zhang, Y., Parmigiani, G. & Johnson, W. E. ComBat-seq: batch effect adjustment for RNA-seq count data. NAR Genom Bioinform 2, lqaa078, doi:10.1093/nargab/lqaa078 (2020).

36 Hart, T. et al. Evaluation and Design of Genome-Wide CRISPR/SpCas9 Knockout Screens. *G3* (*Bethesda*) 7, 2719-2727, doi:10.1534/g3.117.041277 (2017).

37 Li, W. et al. MAGeCK enables robust identification of essential genes from genome-scale CRISPR/Cas9 knockout screens. Genome Biol 15, 554, doi:10.1186/s13059-014-0554-4 (2014).

38 Colic, M. et al. Identifying chemogenetic interactions from CRISPR screens with drugZ. Genome Med 11, 52, doi:10.1186/s13073-019-0665-3 (2019).

39 Hao, Y. et al. Integrated analysis of multimodal single-cell data. Cell 184, 3573–3587 e3529, doi:10.1016/j.cell.2021.04.048 (2021).

40 Ianevski, A., Giri, A. K. & Aittokallio, T. Fully-automated and ultra-fast cell-type identification using specific marker combinations from single-cell transcriptomic data. Nat Commun 13, 1246, doi:10.1038/s41467-022-28803-w (2022).

41 Chen, E. Y. et al. Enrichr: interactive and collaborative HTML5 gene list enrichment analysis tool. BMC Bioinformatics 14, 128, doi:10.1186/1471-2105-14-128 (2013).

42 Kuleshov, M. V. et al. Enrichr: a comprehensive gene set enrichment analysis web server 2016 update. Nucleic Acids Res 44, W90–97, doi:10.1093/nar/gkw377 (2016).

43 Xie, Z. et al. Gene Set Knowledge Discovery with Enrichr. Curr Protoc 1, e90, doi:10.1002/cpz1.90 (2021).

44 Liberzon, A. et al. The Molecular Signatures Database (MSigDB) hallmark gene set collection. Cell Syst 1, 417–425, doi:10.1016/j.cels.2015.12.004 (2015).

45 Knudsen, E. S. et al. Pancreatic cancer cell lines as patient-derived avatars: genetic characterisation and functional utility. Gut 67, 508–520, doi:10.1136/gutjnl-2016-313133 (2018).

46 Zeng, H. et al. Genome-wide CRISPR screening reveals genetic modifiers of mutant EGFR dependence in human NSCLC. Elife 8, doi:10.7554/eLife.50223 (2019).

47 Cooper, J. & Giancotti, F. G. Integrin Signaling in Cancer: Mechanotransduction, Stemness, Epithelial Plasticity, and Therapeutic Resistance. Cancer Cell 35, 347–367, doi:10.1016/j.ccell.2019.01.007 (2019).

48 Cai, X., Wang, K. C. & Meng, Z. Mechanoregulation of YAP and TAZ in Cellular Homeostasis and Disease Progression. Front Cell Dev Biol 9, 673599, doi:10.3389/fcell.2021.673599 (2021).

49 Janes, M. R. et al. Targeting KRAS Mutant Cancers with a Covalent G12C-Specific Inhibitor. Cell 172, 578–589 e517, doi:10.1016/j.cell.2018.01.006 (2018).

50 Fujita-Sato, S. et al. Enhanced MET Translation and Signaling Sustains K-Ras-Driven Proliferation under Anchorage-Independent Growth Conditions. Cancer Res 75, 2851–2862, doi:10.1158/0008-5472.CAN-14-1623 (2015).

51 Beatty, G. L. et al. CD40 agonists alter tumor stroma and show efficacy against pancreatic carcinoma in mice and humans. Science 331, 1612–1616, doi:10.1126/science.1198443 (2011).

52 Dey, P. et al. Oncogenic KRAS-Driven Metabolic Reprogramming in Pancreatic Cancer Cells Utilizes Cytokines from the Tumor Microenvironment. Cancer Discov 10, 608–625, doi:10.1158/2159-8290.CD-19-0297 (2020).

53 Knudsen, E. S. et al. Targeting dual signalling pathways in concert with immune checkpoints for the treatment of pancreatic cancer. Gut 70, 127–138, doi:10.1136/gutjnl-2020-321000 (2021).

54 Kalbasi, A. et al. Uncoupling interferon signaling and antigen presentation to overcome immunotherapy resistance due to JAK1 loss in melanoma. Sci Transl Med 12, doi:10.1126/scitranslmed.abb0152 (2020).

55 Paschen, A., Melero, I. & Ribas, A. Central Role of the Antigen-Presentation and Interferon-γ Pathways in Resistance to Immune Checkpoint Blockade. Annual Review of Cancer Biology 6, 85–102, doi:10.1146/annurev-cancerbio-070220-111016 (2022).

56 Rice, C. M. et al. Tumour-elicited neutrophils engage mitochondrial metabolism to circumvent nutrient limitations and maintain immune suppression. Nat Commun 9, 5099, doi:10.1038/s41467-018-07505-2 (2018).

57 Rogers, T. & DeBerardinis, R. J. Metabolic Plasticity of Neutrophils: Relevance to Pathogen Responses and Cancer. Trends Cancer 7, 700–713, doi:10.1016/j.trecan.2021.04.007 (2021).

58 Uprety, D. & Adjei, A. A. KRAS: From undruggable to a druggable Cancer Target. Cancer Treat Rev 89, 102070, doi:10.1016/j.ctrv.2020.102070 (2020).

59 Kemp, S. B. et al. Efficacy of a Small-Molecule Inhibitor of KrasG12D in Immunocompetent Models of Pancreatic Cancer. Cancer Discov 13, 298–311, doi:10.1158/2159-8290.CD-22-1066 (2023).

60 Schram, A. M. et al. A phase Ib dose-escalation and expansion study of the oral MEK inhibitor pimasertib and PI3K/MTOR inhibitor voxtalisib in patients with advanced solid tumours. Br J Cancer 119, 1471–1476, doi:10.1038/s41416-018-0322-4 (2018).

61 Ko, A. H. et al. A Multicenter, Open-Label Phase II Clinical Trial of Combined MEK plus EGFR Inhibition for Chemotherapy-Refractory Advanced Pancreatic Adenocarcinoma. Clin Cancer Res 22, 61–68, doi:10.1158/1078-0432.CCR-15-0979 (2016).

62 Hallin, J. et al. The KRAS(G12C) Inhibitor MRTX849 Provides Insight toward Therapeutic Susceptibility of KRAS-Mutant Cancers in Mouse Models and Patients. Cancer Discov 10, 54–71, doi:10.1158/2159-8290.CD-19-1167 (2020).

63 Pellinen, T. et al. ITGB1-dependent upregulation of Caveolin-1 switches TGFbeta signalling from tumour-suppressive to oncogenic in prostate cancer. Sci Rep 8, 2338, doi:10.1038/s41598-018-20161-2 (2018).

64 Tiriac, H., Plenker, D., Baker, L. A. & Tuveson, D. A. Organoid models for translational pancreatic cancer research. Curr Opin Genet Dev 54, 7–11, doi:10.1016/j.gde.2019.02.003 (2019).

65 Kumarasamy, V. et al. RB loss determines selective resistance and novel vulnerabilities in ER-positive breast cancer models. Oncogene, doi:10.1038/s41388-022-02362-2 (2022).

66 Li, X. & Wang, C. Y. From bulk, single-cell to spatial RNA sequencing. Int J Oral Sci 13, 36, doi:10.1038/s41368-021-00146-0 (2021).

67 Hu-Lieskovan, S. et al. Improved antitumor activity of immunotherapy with BRAF and MEK inhibitors in BRAF(V600E) melanoma. Sci Transl Med 7, 279ra241, doi:10.1126/scitranslmed.aaa4691 (2015).

68 Jorgovanovic, D., Song, M., Wang, L. & Zhang, Y. Roles of IFN-gamma in tumor progression and regression: a review. Biomark Res 8, 49, doi:10.1186/s40364-020-00228-x (2020).

69 Axelrod, M. L., Cook, R. S., Johnson, D. B. & Balko, J. M. Biological Consequences of MHC-II Expression by Tumor Cells in Cancer. Clin Cancer Res 25, 2392–2402, doi:10.1158/1078-0432.CCR-18-3200 (2019).

