## Supplementary information for "Pharmacologically targeting KRAS^G12D^ in PDAC models: tumor cell intrinsic and extrinsic impact"

**Fig S1.** (A) Live cell imaging to monitor the proliferation of 1222 and 3226 cells following the treatment with MRTX1133 at different concentrations. Error bars were calculated based on mean and SD from triplicates. Experiments were done at two independent times. (B) Differential effect of MRTX1133 and MRTX849 on MiaPaCa2 and UM53 cell lines. Mean and SD were shown from triplicates. Experiments were done at two independent times. (***p<0.0001 as determined by two-way ANOVA). (C) Effect of MRTX1133 on BrdU incorporation on the indicated cell lines following the treatment with different concentrations of MRTX1133. Error bars represent mean and SD from triplicates. (D) Western blotting on the indicated proteins from MiaPaCa-2 cell lines treated with MRTX849 an dMRTX1133.

**Fig. S2** (A) Scatter plot indicating the Normalized count of the sgRNAs from the ASPC1 cells that were selected in DMSO after 11 passages. The essential genes were highlighted. (B) Correlation analysis between the replicates from the CRISPR screen on ASPC1 cells that were selected in the presence and absence of MRTX1133. (C) Effect of FOSL1 KD on the proliferation of ASPC1 cells in combination with MRTX1133. Error bars represent mean and SD from triplicates. (D) Heat map depicting the relative growth rate of ASPC1 cells that were treated with the indicated EGFR inhibitors (100 nM) in combination with DMSO and MRTX1133 (100 nM). (E) Live cell imaging to monitor the growth of ASPC1 cells, treated with MRTX1133 (25 nM) in combination with Gefitinib (500 nM). Mean and SD were shown. (*** p<0.0001 as determined by two-way ANOVA). (F) Effect of MRTX1133 on the depletion of individual guides targeting ITGB1 in ASPC1 cells. (G) Colony formation assay in 3226 and UM53 cell lines, that were treated with MRTX1133 and MRTX849 respectively following the deletion of ITGB1.

**Fig. S3.** (A) Effect of MRTX849 on the growth of UM53 organoids. Mean and SD were shown. (B) Representative images of the UM53 organoids in the presence and absence of MRTX849. (C) Images of tumors excised from AKB6 xenografts that were treated with vehicle and MRTX1133 (30 mg/kg). (D) Pancreas images from a healthy mouse and the genetically modified

KC mouse, bearing the tumor. (E) Effect of MRTX1133 on BrdU incorporation in 4662 cell line. Mean and SD were shown from triplicates. Experiment was at two independent times. (F) Effect of MRTX1133 on the proliferation of AKB6 cell line following the induction of oncogenic $KRAS^{G12D}$ by doxycycline. Error bars represent mean and SD from triplicates. Experiment was repeated at two independent times. (***p<0.0001 as determined by two-way ANOVA). (G) Differential effect of MRTX1133 on the growth of organoid and 2D monolayer derived from 4662 cells. Error bars represent mean and SD from triplicates.

**Fig. S4 :** (A) Single-cell clustering of vehicle and MRTX1133-treated AKB6 tumors to selectively indicate different cellular components. (B) Changes on the indicated cellular population following MRTX1133 treatment. (***p<0.0001 as determined by Fisher Extract test).

**Fig S5.** Seurat heatmap depicting the expression of genes that distinguish different clusters that represent different cellular components.

**Fig. S6:** Pathway analysis to indicate the enrichment of differentially downregulated genes from the PACK+ cells from the tumor tissue derived from AKB6 xenografts following the treatment with MRTX1133. (B) Heat map depicting the downregulated genes across multiple ROIs from the vehicle treated tumor tissues (n=2) and MRTX1133 treated tumor tissues (n=4). Pathway analysis to indicate the enrichment of differentially upregulated genes from the PACK+ cells from the tumor tissue derived from AKB6 xenografts following the treatment with MRTX1133. (D) Heatmap depicting the upregulated genes from ASPC1, HPAF-II and 828 cell lines following the treatment with MRX1133 (100 nM).

**Fig. S7:** (A) Pathway analysis to indicate the enrichment of differentially downregulated genes from the neutrophils population following the treatment with MRTX1133. (B) Seurat feature plots to illustrate the differential expression of representative T-cell markers from the vehicle and MRTX1133-treated groups. (C) Seurat feature plots to illustrate the differential expression of exhaustion markers from the T cell population from the vehicle and MRTX1133-treated groups. (D) Column graph indicating the change in % cell population from CAFs and macrophages between the vehicle and MRTX1133 treated groups. P values were calculated by Fisher Extract test. (E) Seurat feature plots to illustrate the differential expression of representative CAFs markers from the vehicle and MRTX1133-treated groups.

Fig. S1

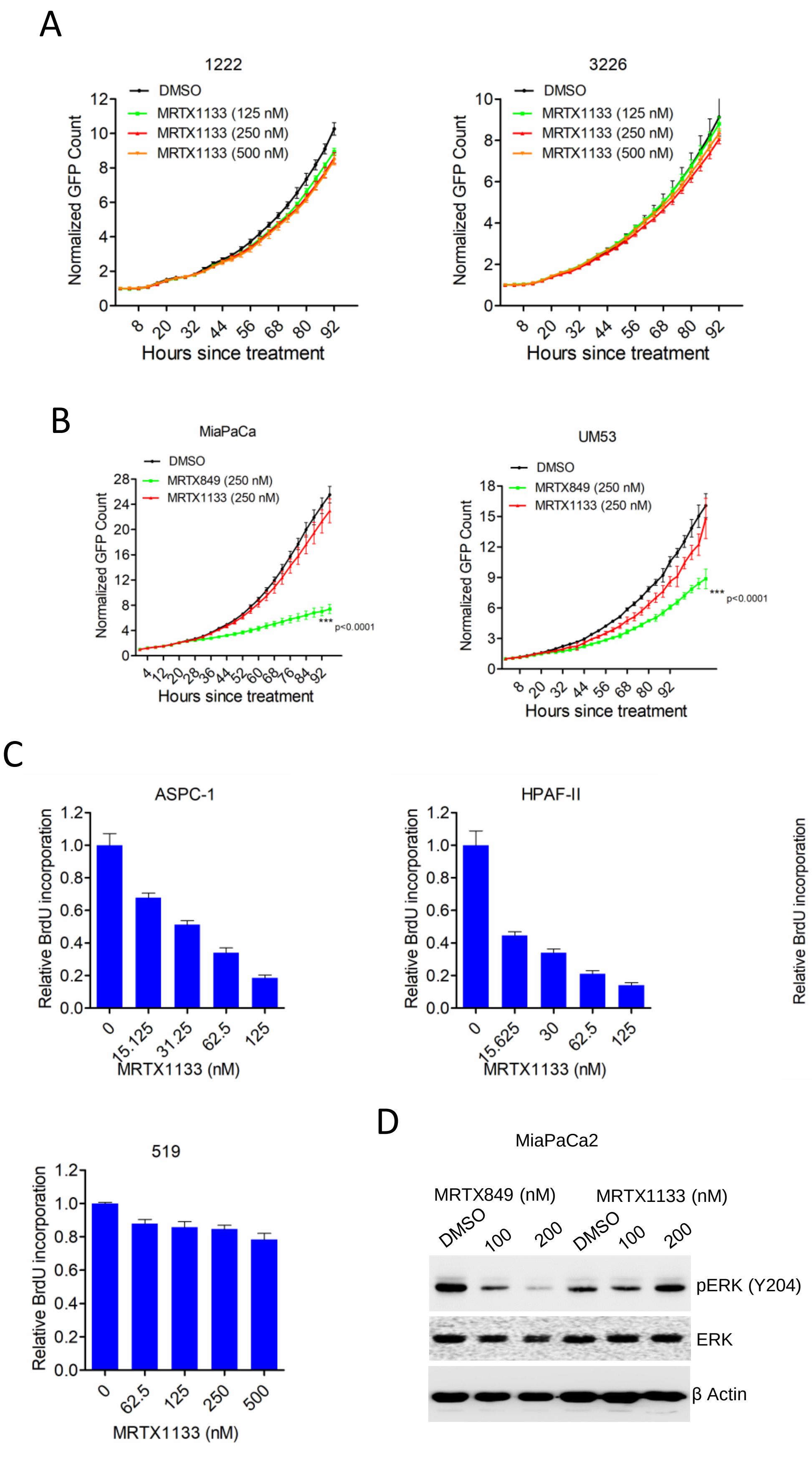
A
1222
DMSO
MRTX1133 (125 nM)
MRTX1133 (250 nM)
MRTX1133 (500 nM)
Normalized GFP Count
Hours since treatment
3226
B
MiaPaCa
DMSO
MRTX849 (250 nM)
MRTX1133 (250 nM)
*** p<0.0001
UM53
C
ASPC-1
HPAF-II
828
519
Relative BrdU incorporation
MRTX1133 (nM)
D
MiaPaCa2
MRTX849 (nM)
MRTX1133 (nM)
DMSO
100
200
pERK (Y204)
ERK
β Actin

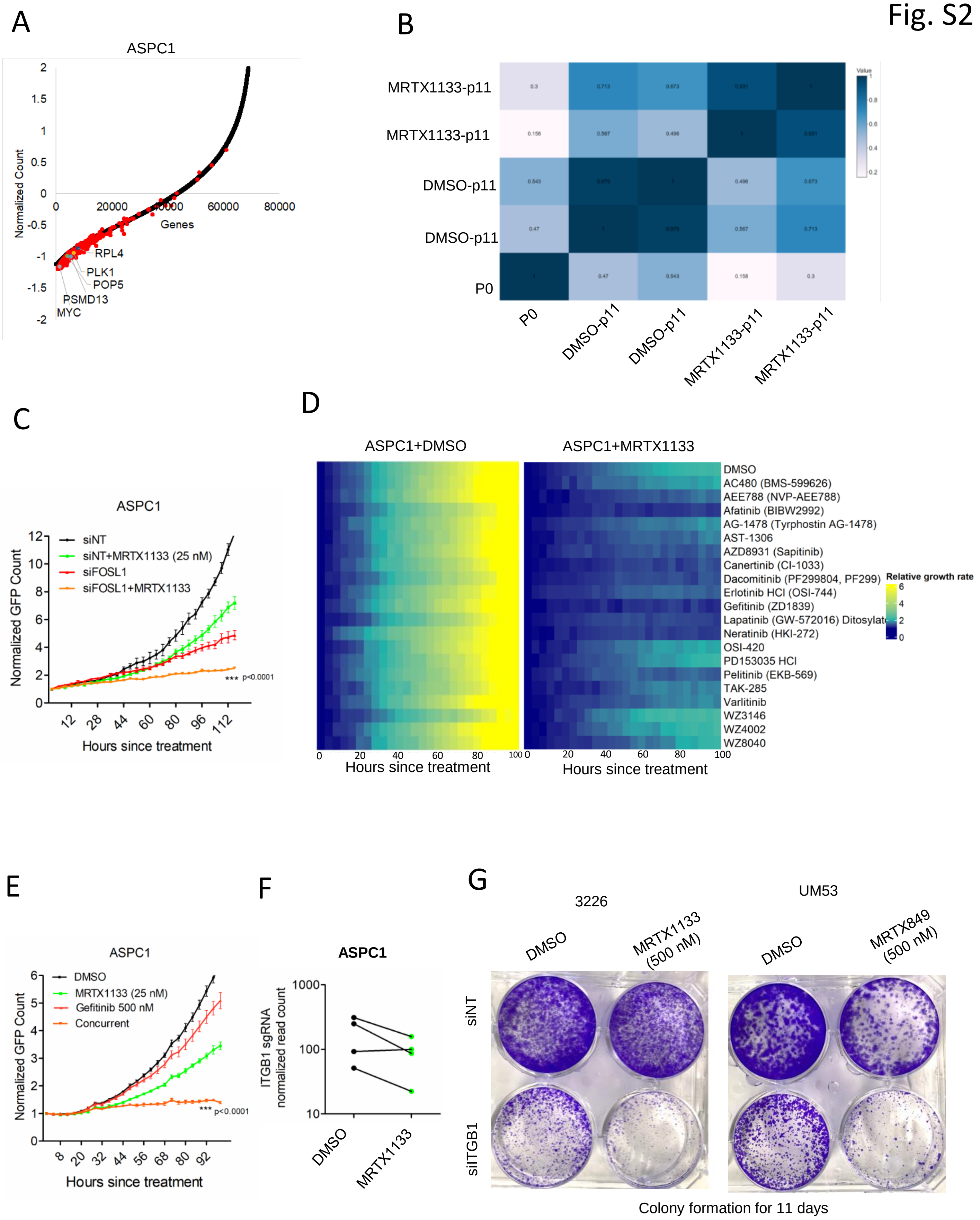
Fig. S2
A
ASPC1
Normalized Count
Genes
RPL4
PLK1
POP5
PSMD13
MYC
B
MRTX1133-p11
MRTX1133-p11
DMSO-p11
DMSO-p11
P0
Value
C
ASPC1
siNT
siNT+MRTX1133 (25 nM)
siFOSL1
siFOSL1+MRTX1133
Normalized GFP Count
*** p<0.0001
Hours since treatment
D
ASPC1+DMSO
ASPC1+MRTX1133
DMSO
AC480 (BMS-599626)
AEE788 (NVP-AEE788)
Afatinib (BIBW2992)
AG-1478 (Tyrphostin AG-1478)
AST-1306
AZD8931 (Sapitinib)
Canertinib (CI-1033)
Dacomitinib (PF299804, PF299)
Erlotinib HCl (OSI-744)
Gefitinib (ZD1839)
Lapatinib (GW-572016) Ditosylate
Neratinib (HKI-272)
OSI-420
PD153035 HCl
Pelitinib (EKB-569)
TAK-285
Varlitinib
WZ3146
WZ4002
WZ8040
Relative growth rate
Hours since treatment
Hours since treatment
E
ASPC1
DMSO
MRTX1133 (25 nM)
Gefitinib 500 nM
Concurrent
Normalized GFP Count
*** p<0.0001
Hours since treatment
F
ASPC1
ITGB1 sgRNA normalized read count
DMSO
MRTX1133
G
3226
UM53
DMSO
MRTX1133 (500 nM)
DMSO
MRTX849 (500 nM)
siNT
siITGB1
Colony formation for 11 days

Fig. S3

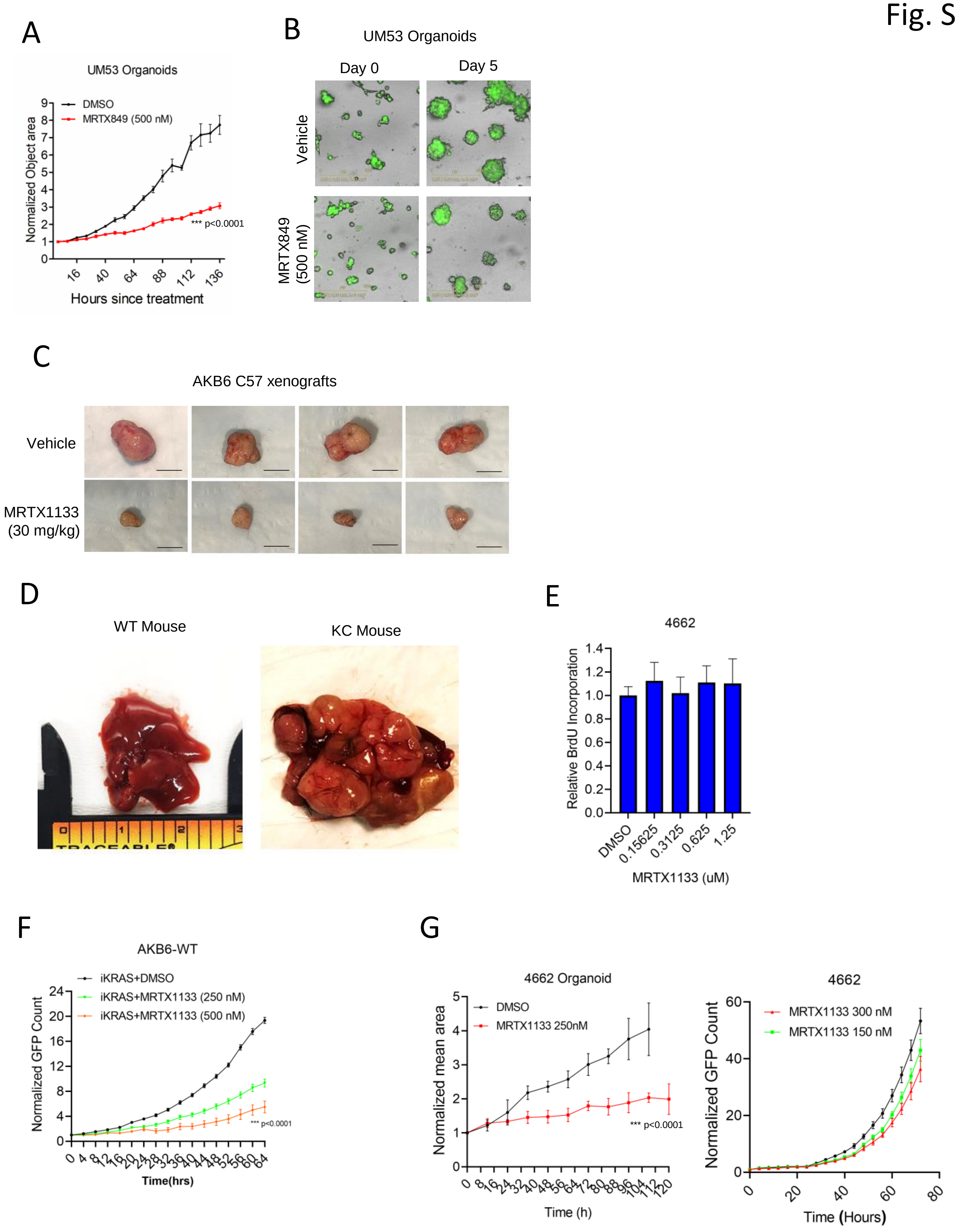
A
UM53 Organoids
DMSO
MRTX849 (500 nM)
Normalized Object area
*** p<0.0001
Hours since treatment
B
UM53 Organoids
Day 0
Day 5
Vehicle
MRTX849 (500 nM)
C
AKB6 C57 xenografts
Vehicle
MRTX1133 (30 mg/kg)
D
WT Mouse
KC Mouse
E
4662
Relative BrdU Incorporation
DMSO
0.15625
0.3125
0.625
1.25
MRTX1133 (uM)
F
AKB6-WT
iKRAS+DMSO
iKRAS+MRTX1133 (250 nM)
iKRAS+MRTX1133 (500 nM)
Normalized GFP Count
*** p<0.0001
Time(hrs)
G
4662 Organoid
DMSO
MRTX1133 250nM
Normalized mean area
*** p<0.0001
Time (h)
4662
MRTX1133 300 nM
MRTX1133 150 nM
Normalized GFP Count
Time (Hours)

### AKB6 Xenografts

A

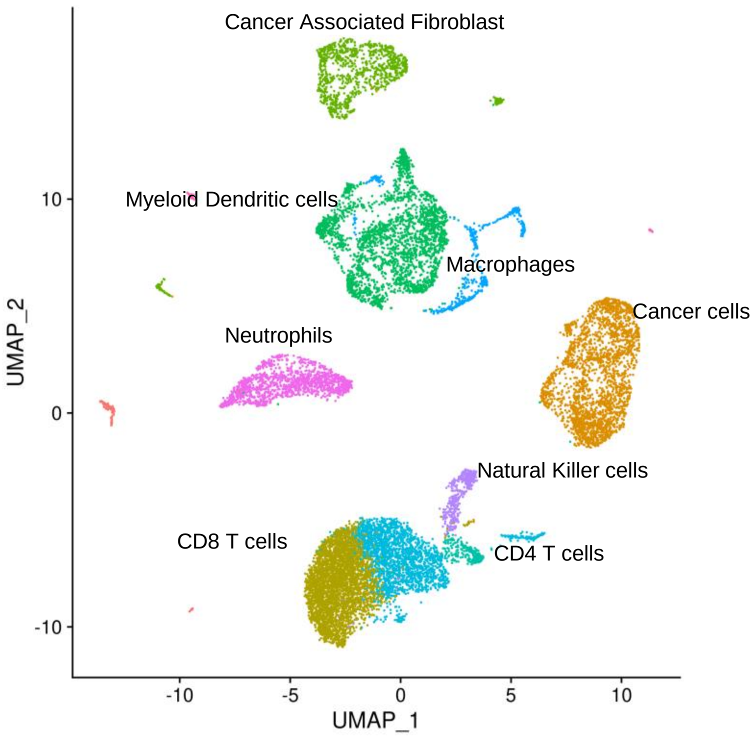


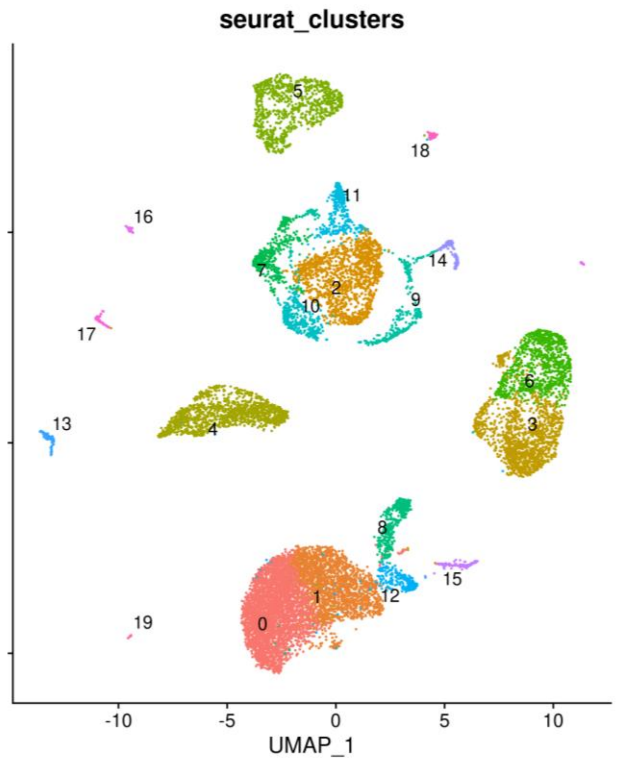


B

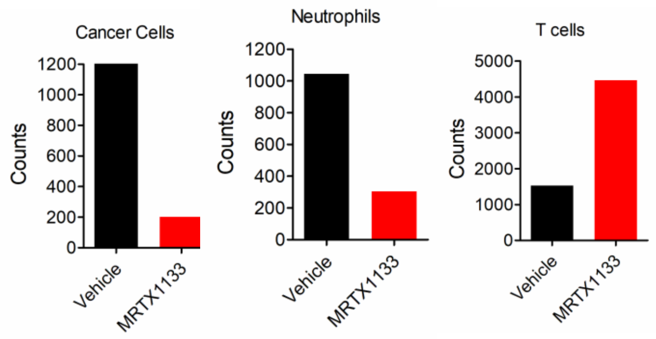

Fig. S5

A

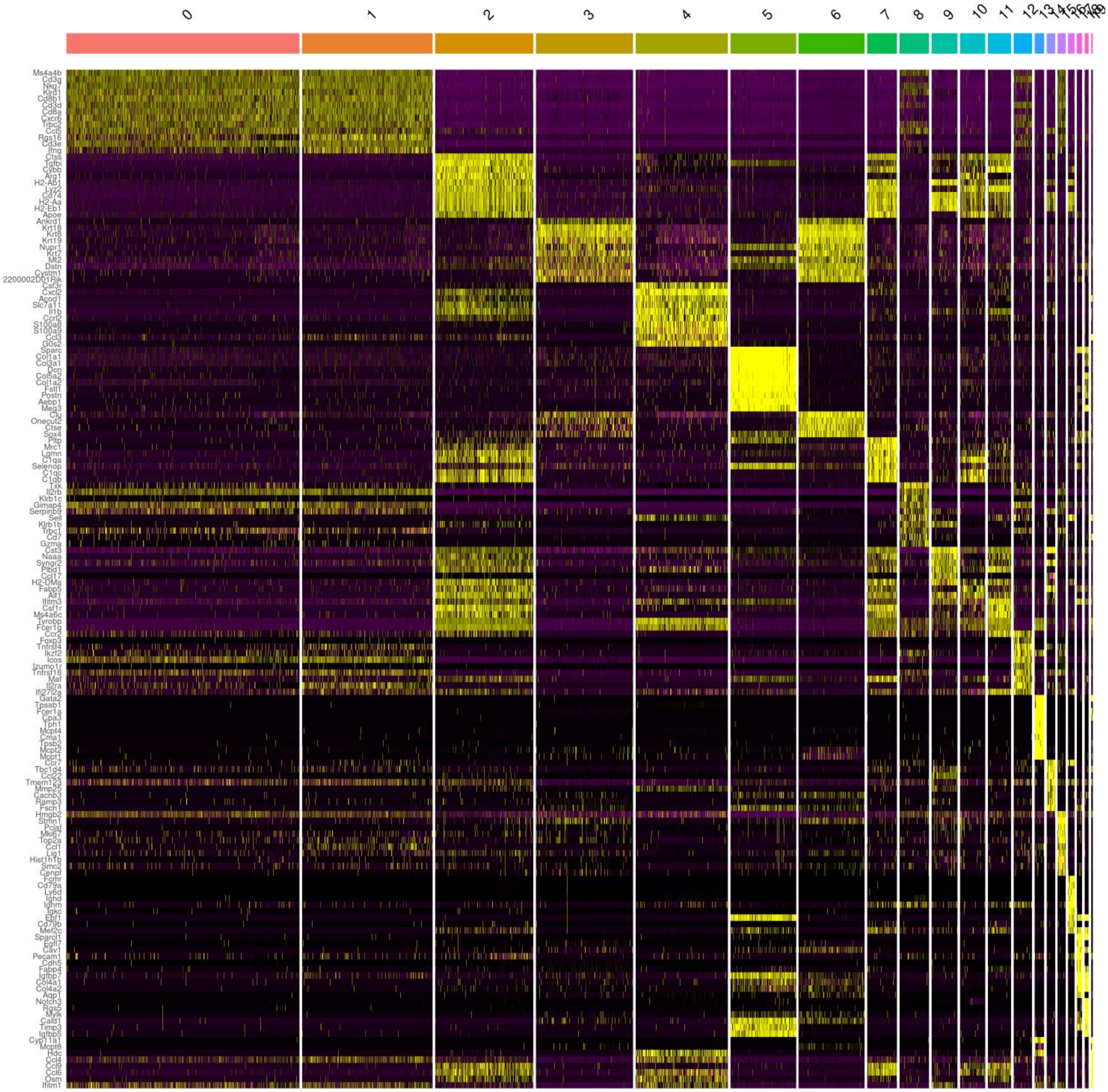
0
1
2
3
4
5
6
7
8
9
10
11
12
Ms4a4b
Cd3g
Nkg7
Klrd1
Cd8b1
Cd3d
Cd8a
Cxcr6
Trbc2
Ccl5
Rgs16
Cd3e
Ifng
Ctss
Tgfbi
Cybb
Arg1
H2-Ab1
Lyz2
Cd74
H2-Aa
H2-Eb1
Apoe
Ankrd1
Krt18
Krt8
Krt19
Nupr1
Krt7
Mt2
Dstn
Cystm1
2200002D01Rik
Csf3r
Cxcl2
Acod1
Slc7a11
Il1b
Ccrl2
S100a8
S100a9
Ccl3
G0s2
Sparc
Col1a1
Col3a1
Dcn
Col5a2
Col1a2
Fstl1
Postn
Aebp1
Meg3
Clu
Onecut2
Ctse
Sox4
Pltp
Mrc1
Lgmn
C1qa
Selenop
C1qc
C1qb
Txk
Il2rb
Klrb1c
Gimap4
Serpinb9
Sell
Klrb1b
Trbc1
Cd7
Gzma
Cst3
Naaa
Syngr2
Plbd1
Ccl17
H2-DMa
Fabp5
Aif1
Ifitm3
Csf1r
Ms4a6c
Tyrobp
Fcer1g
Ccr2
Foxp3
Tnfrsf4
Ikzf2
Icos
Izumo1r
Tnfrsf18
Maf
Il2ra
Ifi27l2a
Gata2
Tpsab1
Fcer1a
Cpa3
Tph1
Mcpt4
Cma1
Tpsb2
Mcpt2
Mcpt1
Ccr7
Tbc1d4
Ccl22
Tmem123
Mmp25
Cacnb3
Ramp3
Fscn1
Hmgb2
Stmn1
Pclaf
Mki67
Top2a
Ccl1
Lig1
Hist1h1b
Smc2
Cenpf
Fcmr
Cd79a
Ly6d
Ighd
Ighm
Igkc
Ebf1
Cd79b
Mef2c
Sparcl1
Egfl7
Cav1
Pecam1
Cdh5
Fabp4
Igfbp7
Col4a1
Col4a2
Aqp1
Notch3
Rgs5
Mylk
Cald1
Timp3
Igfbp5
Cyp11a1
Mcpt8
Hdc
Ccl4
Ccl9
Ccl6
Osm
Ifitm1

Fig. S6

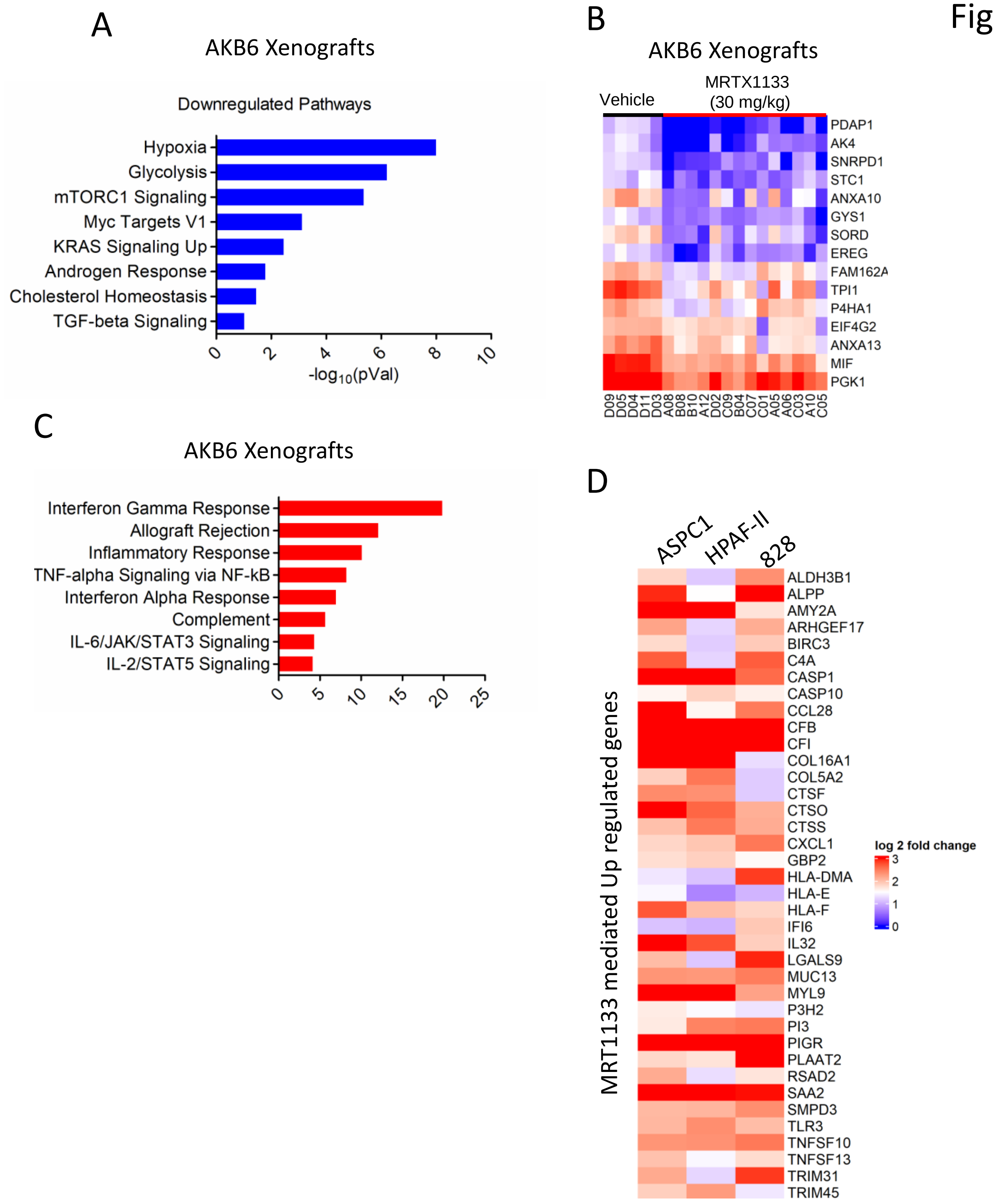
A
AKB6 Xenografts
Downregulated Pathways
Hypoxia
Glycolysis
mTORC1 Signaling
Myc Targets V1
KRAS Signaling Up
Androgen Response
Cholesterol Homeostasis
TGF-beta Signaling
0
2
4
6
8
10
-log10(pVal)
B
AKB6 Xenografts
Vehicle
MRTX1133 (30 mg/kg)
PDAP1
AK4
SNRPD1
STC1
ANXA10
GYS1
SORD
EREG
FAM162A
TPI1
P4HA1
EIF4G2
ANXA13
MIF
PGK1
D09
D05
D04
D11
D03
A08
B08
B10
A12
D02
C09
B04
C07
C01
A05
A06
C03
A10
C05
C
AKB6 Xenografts
Interferon Gamma Response
Allograft Rejection
Inflammatory Response
TNF-alpha Signaling via NF-kB
Interferon Alpha Response
Complement
IL-6/JAK/STAT3 Signaling
IL-2/STAT5 Signaling
0
5
10
15
20
25
D
ASPC1
HPAF-II
828
MRT1133 mediated Up regulated genes
ALDH3B1
ALPP
AMY2A
ARHGEF17
BIRC3
C4A
CASP1
CASP10
CCL28
CFB
CFI
COL16A1
COL5A2
CTSF
CTSO
CTSS
CXCL1
GBP2
HLA-DMA
HLA-E
HLA-F
IFI6
IL32
LGALS9
MUC13
MYL9
P3H2
PI3
PIGR
PLAAT2
RSAD2
SAA2
SMPD3
TLR3
TNFSF10
TNFSF13
TRIM31
TRIM45
log 2 fold change
3
2
1
0

Fig. S7

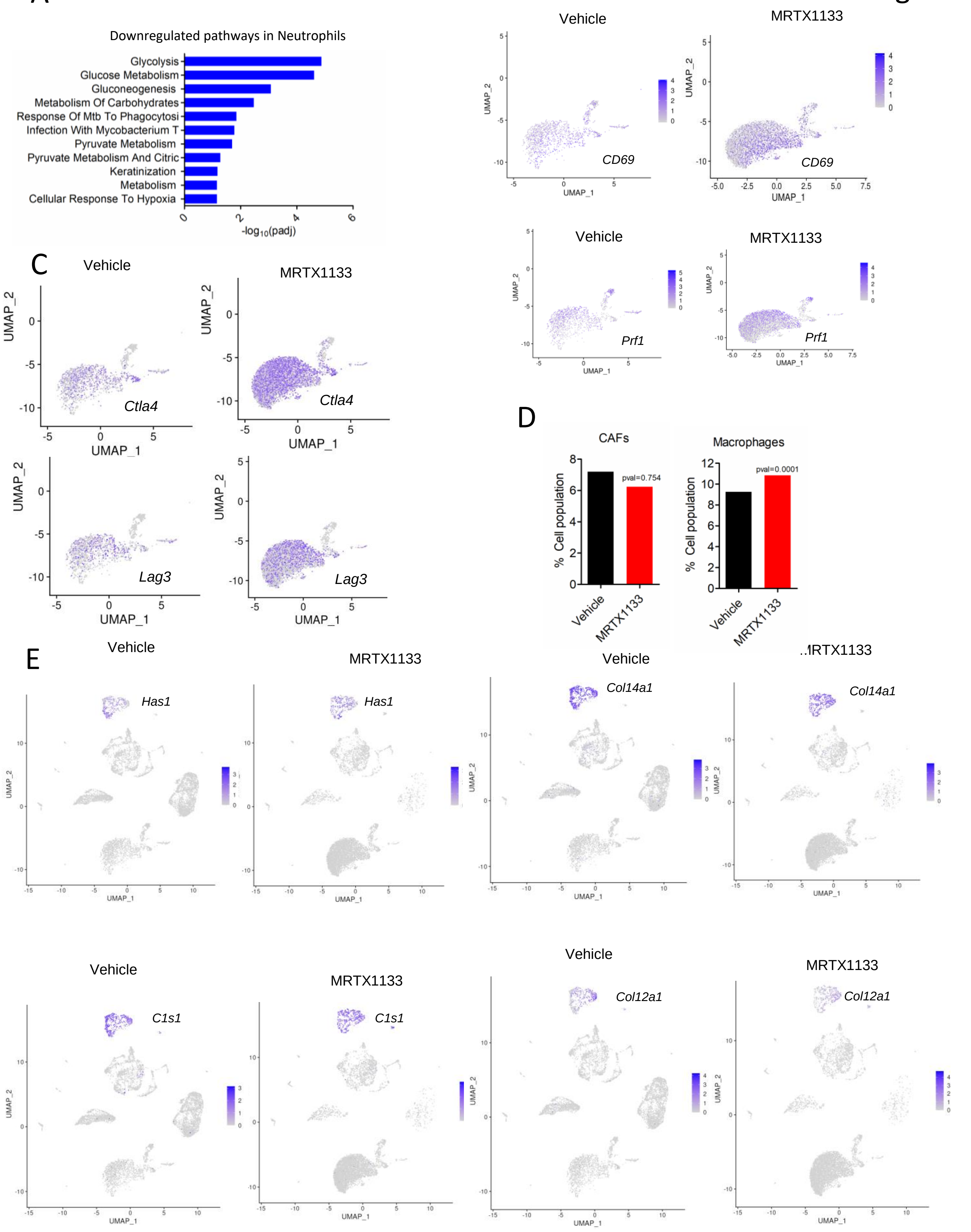
A
Downregulated pathways in Neutrophils
Glycolysis
Glucose Metabolism
Gluconeogenesis
Metabolism Of Carbohydrates
Response Of Mtb To Phagocytosi
Infection With Mycobacterium T
Pyruvate Metabolism
Pyruvate Metabolism And Citric
Keratinization
Metabolism
Cellular Response To Hypoxia
-log10(padj)
B
Vehicle
MRTX1133
CD69
Prf1
UMAP_1
UMAP_2
C
Vehicle
MRTX1133
Ctla4
Lag3
UMAP_1
UMAP_2
D
CAFs
Macrophages
% Cell population
pval=0.754
pval=0.0001
Vehicle
MRTX1133
E
Vehicle
MRTX1133
Has1
Col14a1
C1s1
Col12a1
UMAP_1
UMAP_2
